# zol & fai: large-scale targeted detection and evolutionary investigation of gene clusters

**DOI:** 10.1101/2023.06.07.544063

**Authors:** Rauf Salamzade, Patricia Q. Tran, Cody Martin, Abigail L. Manson, Michael S. Gilmore, Ashlee M. Earl, Karthik Anantharaman, Lindsay R. Kalan

## Abstract

Many universally and conditionally important genes are genomically aggregated within clusters. Here, we introduce fai and zol, which together enable large-scale comparative analysis of different types of gene clusters and mobile-genetic elements (MGEs), such as biosynthetic gene clusters (BGCs) or viruses. Fundamentally, they overcome a current bottleneck to reliably perform comprehensive orthology inference at large scale across broad taxonomic contexts and thousands of genomes. First, fai allows the identification of orthologous instances of a query gene cluster of interest amongst a database of target genomes. Subsequently, zol enables reliable, context-specific inference of ortholog groups for individual protein-encoding genes across gene cluster instances. In addition, zol performs functional annotation and computes a variety of evolutionary statistics for each inferred ortholog group. Importantly, in comparison to tools for visual exploration of homologous relationships between gene clusters, zol can scale to thousands of gene cluster instances and produce detailed reports that are easy to digest. To showcase fai and zol, we apply them for: (i) longitudinal tracking of a virus in metagenomes, (ii) discovering novel population-level genetic insights of two common BGCs in the fungal species *Aspergillus flavus*, and (iii) uncovering large-scale evolutionary trends of a virulence-associated gene cluster across thousands of genomes from a diverse bacterial genus.

## Background

*De novo* ortholog grouping typically involves searching for reciprocal best hits of proteins between pairs of genomes, indicative of orthology, and subsequently clustering pairs of inferred orthologs and in-paralogs across multiple genomes^1–4^. Initial methods for orthology inference were designed to be able to identify orthologs between distinct species but limited in the number of genomes they could process^1–3^. This limitation is largely due to the all-vs-all alignment of proteomes, core to most methods for *de novo* ortholog grouping, which is an *O*(*n*^2^) operation and a major computational bottleneck. Approaches to overcome this procedure include limiting proteome comparisons by using a guiding-phylogeny^5,6^, adapting alignment searching parameters and heuristics to further boost speeds^7,8^, or preliminary aggressive clustering of proteins into coarse homolog groups^9^. Recently, graph-based and iterative-clustering approaches have also allowed vast scalability to thousands of bacterial genomes, but are primarily designed for application to a single species^10–13^.

Available orthology inference methods struggle to infer ortholog groups across large datasets of taxonomically diverse genomes, potentially representing thousands of species, such as a set of metagenome-assembled genomes (MAGs) related to a common microbiome. While multiple methods exist to identify instances of previously established ortholog groups within the predicted proteome of a metagenome^14–17^, these are unable to account for proteins not represented in their database. Recently, independent advancements in methods to collapse large protein sets based on sequence similarity have enabled rapid clustering of millions of sequences^18–20^. These approaches have even been used on massive protein datasets gathered from across multiple metagenomic datasets^21^; however, more resolute delineation of functionally analogous ortholog groups across thousands of genomes from multiple species remains difficult to perform *de novo*.

Of relevance, within bacterial genomes, genes are often co-located within smaller, discrete, multi-gene units, which we will broadly refer to as gene clusters. Examples of gene clusters include operons^22,23^, phages^24^, metabolic gene clusters^25^, biosynthetic gene clusters (BGCs)^26–29^, and pathogenicity islands^30,31^. Although less common, eukaryotic genomes can also contain genes aggregated within discrete clusters^32–34^. Sometimes gene clusters are highly conserved, encoding for products essential to the survival of the organism^35^. In other cases, a single gene cluster can exhibit variability in gene carriage and order across different strains or species^36–38^. This is often the case for BGCs encoding specialized metabolites or virulence-associated gene clusters, where evolution of gene content and sequence divergence can influence fitness and contribute to adaptation within a changing ecosystem^39–41^.

Syntenic conservation has been used to assist *de novo* identification of homologous instances of a gene cluster of interest in diverse target genomes^42–45^. Homologous gene cluster instances can then be comprehensively investigated to delineate homolog or ortholog groups of the proteins found across them^44,46^. While such targeted approaches can alleviate time and computational resources by avoiding more comprehensive identification of orthologs at genome-wide scales, currently available methods are mostly designed for specific types of gene clusters, such as BGCs^42,44,45^. Many of the software implementing such approaches also do not provide support for uniform annotation of coding sequences in target genomes, which can decrease sensitivity for gene cluster detection. In addition, most methods do not account for gene cluster paralogy, which has been observed for BGCs in bacterial^38^ and fungal genomes^33^, or provide specialized capabilities for finding gene clusters across fragmented genomes or metagenomic assemblies^38^.

Following identification of homologous gene clusters in target genomes, software to understand the evolutionary relationships between gene cluster instances and infer protein ortholog groups have largely applied coarse protein clustering and aimed to provide visualization based exploration to users^44,46–48^. Visual assessment of related gene clusters and manual refinement of ortholog groups work well at smaller scales but become impractical when dealing with hundreds to thousands of gene cluster instances. Scalability challenges are due to both computational costs needed to render visuals as well as the figures becoming convoluted and difficult to interpret. An effective solution to ease the identification of evolutionary trends amongst homologous gene clusters is to first identify ortholog groups^44^ and present information pertaining to their conservation and sequence divergence within tabular reports^10,38^. Such tabular reports scale by the number of unique ortholog groups and can be organized by their consensus order along gene cluster instances. We recently introduced construction of such reports in a software suite for exploring microdiversity amongst homologous BGCs from a single taxon^38^; however, the functionality was difficult to use outside of the suite and reliant on orthologous relationships between proteins of gene clusters being known in advance.

Here, we introduce the zol suite, providing functionalities for gene cluster detection and subsequent inference and investigation of protein ortholog groups across homologous gene clusters. The versatility and scalability of these programs is demonstrated through application to three types of gene clusters within different genomic contexts including a virus within environmental metagenomes, fungal secondary metabolite encoding biosynthetic gene clusters, and a conserved polysaccharide antigen locus from the diverse bacterial genus of *Enterococcus*.

## Results

### fai and zol allow for the rapid inference of gene cluster orthologs across diverse genomes

The zol suite consists of three major programs: prepTG (*<u>prep</u>are target <u>g</u>enomes*), fai (*find <u>a</u>dditional instances*), and zol (*<u>z</u>oom <u>o</u>n locus*) (**Figure 1A**). First, prepTG and fai can be run to process a set of target genomes and rapidly search for a query gene cluster within them, respectively. Afterwards, zol can perform reliable and efficient context-limited inference of ortholog groups across homologous gene cluster instances identified using a flexible InParanoid-type algorithm^3^. For each ortholog group, zol will further compute evolutionary statistics, such as Tajima’s D^49^, and functional annotations, using several, diverse databases suitable for a variety of gene clusters, including those specific to phages^50^, virulence elements^51^, and BGCs^52^. Ultimately, zol will summarize data in a table report where each row corresponds to a distinct ortholog group. This report is automatically color formatted and provided as an XLSX spreadsheet to allow for easy interpretation of the data, which can span thousands of gene cluster instances.

**Figure 1:**
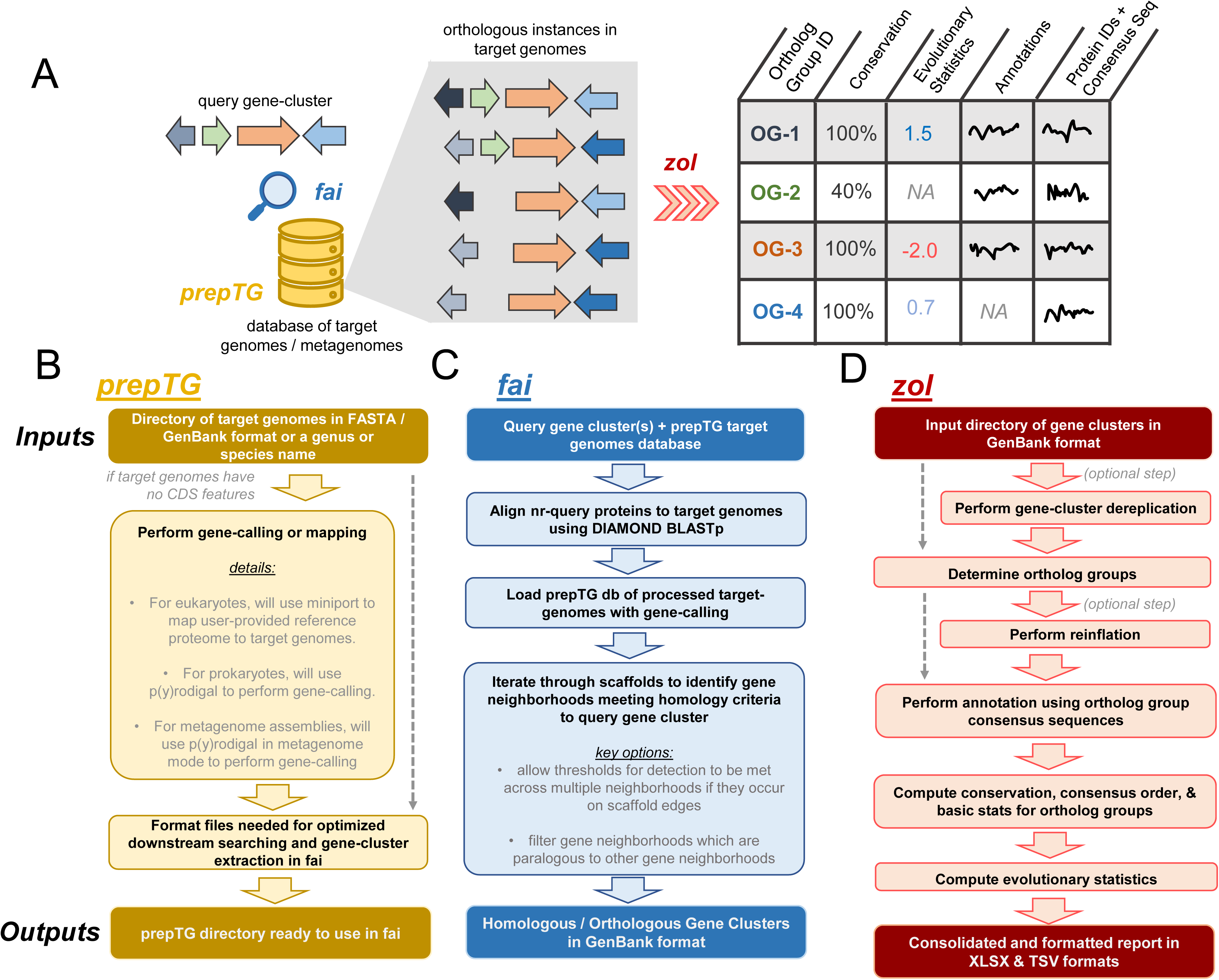
Overviews of fai and zol. **A**) A cartoon schematic of how prepTG, fai, and zol are integrated to perform evolutionary investigations by searching for gene-clusters. Certain statistics in the zol report will not be calculated if not enough instances of an ortholog group are identified, resulting in non-available (NA) values being reported. Squiggles correspond to arbitrary text pertaining to functional annotation information, etc. **B**) An overview of the prepTG, **C**) fai, and **D**) zol algorithms and workflows. Inputs and outputs for the programs are indicated with bolder coloring.

To promote consistency in gene calling across target genomes, we have incorporated computationally light-weight dependencies for *de novo* gene prediction in bacterial genomes^53,54^ and protein-mapping in eukaryotic genomes^55^ within prepTG, to prepare and format target genomes for optimized gene cluster searching in fai (**Figure 1B**). prepTG also aims to provide a convenient interface to transform genomic or metagenomic datasets into a format ready for searching using fai. Options are available to download pre-built databases of distinct representative genomes for 18 commonly studied bacterial taxa^56^ or to build comprehensive databases for any genus or species in the latest release of the Genome Taxonomy Database (GTDB)^57^.

fai features two key features which are absent in most existing methods for gene cluster detection (**Figure 1C; Table S1; Supplementary Text**). First, it has an option to automatically filter secondary instances of query gene clusters identified in target genomes, removing potentially paralogous gene clusters from downstream investigations. Second, fai implements a mode for searching for gene clusters in draft quality genomes, MAGs, or unbinned metagenomic assemblies, where gene clusters might be fragmented across multiple scaffolds. When this mode is activated, fai relaxes requirements for reporting a gene cluster as present in a genome or metagenome if multiple homologous gene cluster regions are identified near scaffold edges in a target genome and instead assesses whether reporting criteria are met in unison across such instances (**Figure S1**). Similar to prepTG, fai also aims to provide convenience for users and can accept query gene clusters in different formats to ease searching for gene clusters and genomic islands cataloged in databases such as ICEberg^58^, MIBiG^52^, or IslandViewer^59^. Query gene clusters can be provided as a coordinate along a reference genome, in GenBank format, or as a set of proteins in FASTA format. In addition, to simplify conservation and novelty assessment of a single isolate’s BGCs, phages, and plasmids relative to other genomes from the same genus or species, specialized wrapper programs of fai are also provided within the zol suite (**Figure S2**).

zol will infer ortholog groups for proteins across homologous gene clusters and then construct a tabular report with information on conservation, evolutionary trends, and annotation for each individual ortholog group (**Figure 1D**). To make annotated reports generated by zol more comprehensive for different types of gene clusters, several databases have been included, such as VOGs^50^, VFDB^51^, ISFinder^60^, and CARD^61^. In addition, zol incorporates HyPhy^62^ as a dependency and calculates various evolutionary statistics. Ultimately, beyond high-throughput inference of ortholog groups across diverse genomic datasets, the rich tabular report produced by zol provides complementary information to figures generated by comparative visualization software such as clinker^46^, CORASON^44^, gggenomes^63^, and Easyfig^64^.

A key feature in zol is the ability to dereplicate gene clusters directly using skani^65^, which was recently shown to be more reliable at estimating average nucleotide identity (ANI) between genomes of variable contiguity relative to comparative methods. Dereplication can allow for more appropriate inference of evolutionary statistics to overcome availability or sampling biases in genomic databases^66^. It can also be used to subset distinct representative gene cluster instances to make investigation using visualization software more tractable. Another important ability of zol is a mode where users can provide a handful of known instances for a gene cluster to estimate optimal parameters to search for additional instances of the gene cluster using fai. We applied this functionality of zol on sets of homologous BGCs and phages to determine distributions for search parameters in fai which users could consult as priors (**Figure S3; Supplementary Text**).

Finally, zol allows for comparative investigations of gene clusters based on taxonomic or ecological groupings^67–69^. For instance, users can designate a subset of gene clusters as belonging to a specific population to allow zol to calculate ortholog group conservation across just the focal set of gene clusters. In addition, zol will compute the fixation index^70^, F_ST_, for each ortholog group to assess gene flow between the focal and complementary sets of gene clusters.

### Longitudinal tracking of a virus within lake metagenomic assemblies

Metagenomic datasets represent a large reservoir of underexplored sequence space^71,72^. To demonstrate the ability of the zol suite to identify and investigate gene clusters in metagenomes, we applied it to track a virus in a longitudinal metagenomic dataset profiling a lake’s microbiome over space and time^73^.

We first identified large (*≥*20kb) viruses, that were also predicted to represent circular molecules, across a subset of the metagenomic assemblies corresponding to the earliest sampling date^74^. Afterwards, clustering based on the sequence and syntenic similarity of protein domains led to the identification of a ∼36kb highly conserved virus in two of the metagenomes sampled from lower lake depths.

All 16 metagenomic assemblies, spanning five distinct sampling timepoints and four distinct sampling depths, were processed through prepTG to identify coding sequences and construct a database ready to search for gene clusters using fai. GenBank files with coding sequence annotations for metagenomic assemblies generated by prepTG, amassing 27 Gb total in size, were further provided as input for cblaster makedb, which serves a similar role to prepTG in the cblaster suite to format genomic data for downstream gene cluster searches. However, cblaster makedb does not feature the ability to perform *de novo* gene-calling for either genomes or metagenomes and is not designed to accommodate the size of metagenomic assemblies. During database construction, cblaster makedb required around 30 Gb of memory, while prepTG needed less than 3 Gb of memory (**Figure S4A**).

Next, fai was used to perform a rapid, targeted search for this ∼36 kb *Caudovirales* virus across the full set of 16 metagenomes to identify additional instances of the virus. fai completed its search of the metagenomes, featuring >20 million proteins and 10.7 million contigs, in less than four minutes using 20 threads, performing similarly to cblaster, run using similar settings as fai (**Figure S4B**). Of the 16 total metagenomes, the virus was found in ten metagenomes, including all nine metagenomes surveying anoxic conditions (*p*<0.001; one-sided Fisher’s exact test; **Figure 2A**). This is concordant with inferences for the host for the virus being *Rhodoferax*, which are purple bacterium featuring species classified as anaerobic photoheterotrophs^73,75,76^. In addition, *Rhodoferax* classified MAGs from the metagenomic dataset were exclusively obtained from anoxic conditions^73^. To investigate how the gene repertoire of the virus evolved over time, we next applied zol. zol-based analysis revealed that 45 (72.6%) of the 62 total distinct ortholog groups were core to all instances of the virus across ten metagenomes with most completely conserved in sequence over the course of 2.5 months (**Figure 2B; Table S2**). Furthermore, 15 of the 62 ortholog groups were not observed in the query viruses from the earliest sampling date, suggesting the potential acquisition or duplication of genes in the virus during the span of sampling at the lake.

**Figure 2:**
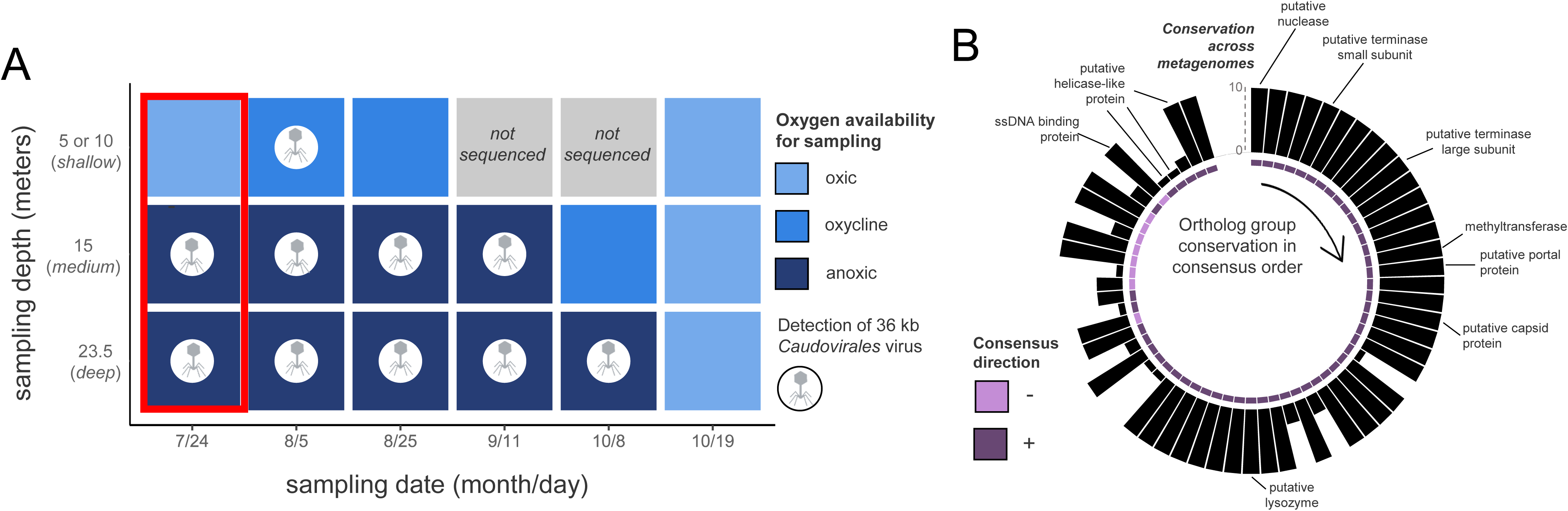
Targeted viral detection in metagenomes using fai. **A**) Total metagenomes from a single site in Lake Mendota across multiple depths and timepoints from Tran et al. 2023 were investigated using fai for the presence of a virus found in two of the three earliest microbiome samplings (red box). The presence of the virus is indicated by a virus icon. Metagenome samples are colored according to whether they corresponded to oxic, oxycline, or anoxic. The most shallow sampling depths varied for different dates and consolidated as a single row corresponding to a sampling depth of either 5 or 10 meters. **D**) The pangenome of the virus is shown based on the consensus order and directionality of coding sequences inferred by zol. Bar heights correspond to the conservation of the ortholog groups across the ten metagenomes the virus was detected in. BioRender was used in generation of this figure.

### Investigating population-level and species-wide evolutionary trends of BGCs in the eukaryotic species *Aspergillus flavus*

Low sensitivity for gene cluster detection in eukaryotic genome assemblies can arise from their incompleteness, leading to gene clusters being fragmented across multiple scaffolds^77,78^, as well as challenges in *ab initio* gene prediction due to alternative splicing^79,80^. Therefore, many gene cluster detection software are either specific for bacterial genomes or require coding sequence annotations for eukaryotic genomes to be provided by the user. To overcome such challenges to user application, we integrated miniprot^55^ into prepTG which allows for mapping high-quality protein annotations from a reference genome to the remainder of the genomes available for a species or genus. We showcase the ability of prepTG and fai to simplify the reliable identification of gene clusters in eukaryotic genomes by using them to find instances of two BGCs across genomes belonging to the fungal species *Aspergillus flavus*.

The genus of *Aspergillus* is a source of several natural products, including aflatoxins, a common and economically impactful contaminant of food^81^. The genus also contains species that are model organisms for studying fungal secondary metabolism^34,82,83^. Examination of the secondary metabolome of *A. flavus* has revealed that different clades or populations can exhibit variability in their metabolite production despite high conservation of core BGC genes encoding enzymes for synthesis of these metabolites^37,84^. For instance, population B *A. flavus* were identified as producing a greater abundance of the insecticide leporin B relative to populations A and C^37,85^. We showcase zol’s ability to aid comparative analysis of gene clusters from different populations through application to the leporin BGC. We further show how zol can detect variation in sequence conservation for different genes from the aflatoxin BGC and be inclusive of genes present in target genome annotations but missing in the query gene cluster, allowing for comprehensive profiling of BGC auxiliary content.

Based on read alignment to a reference genome, the leporin cluster was recently identified to be a core component of the *A. flavus* genome^37^. However, a restricting factor in the direct prediction of gene clusters in *A. flavus* assemblies is the lack of gene annotations, with only 11 (5.1%) of 216 genomes from the species in NCBI’s GenBank database having coding sequence predictions (**Figure 3A**). Therefore, we mapped high-quality protein predictions for a reference *A. flavus* genome^86^ to the remainder of the 216 genomes available for the species. Running fai in “draft mode” led to the identification of the leporin BGC within 212 (98.1%) assemblies, consistent with the prior read mapping-based investigation suggesting that the BGC was core to the species^37^. In comparison, the CAGECAT server^87^, which runs cblaster^45^, was limited to genomes with protein coding annotations available on NCBI and thus unable to assess the remaining 205 genomes for the presence of the leporin BGC (**Figure 3B**). We also investigated the ability of non-targeted approaches for BGC detection to identify the leporin BGC by applying antiSMASH followed by BiG-SCAPE for clustering related BGCs and matching them to characterized BGCs in the MIBiG database. When this approach was applied using GenBank files prepared by prepTG, the gene cluster clan corresponding containing the leporin BGC was found in all *A. flavus* genomes provided as input. However, when antiSMASH was run using *de novo* gene prediction in antiSMASH based on GlimmerHMM^88^ with *Cryptococcus* gene annotation models, recovery of the leporin BGC was limited (**Figure 3B**).

**Figure 3:**
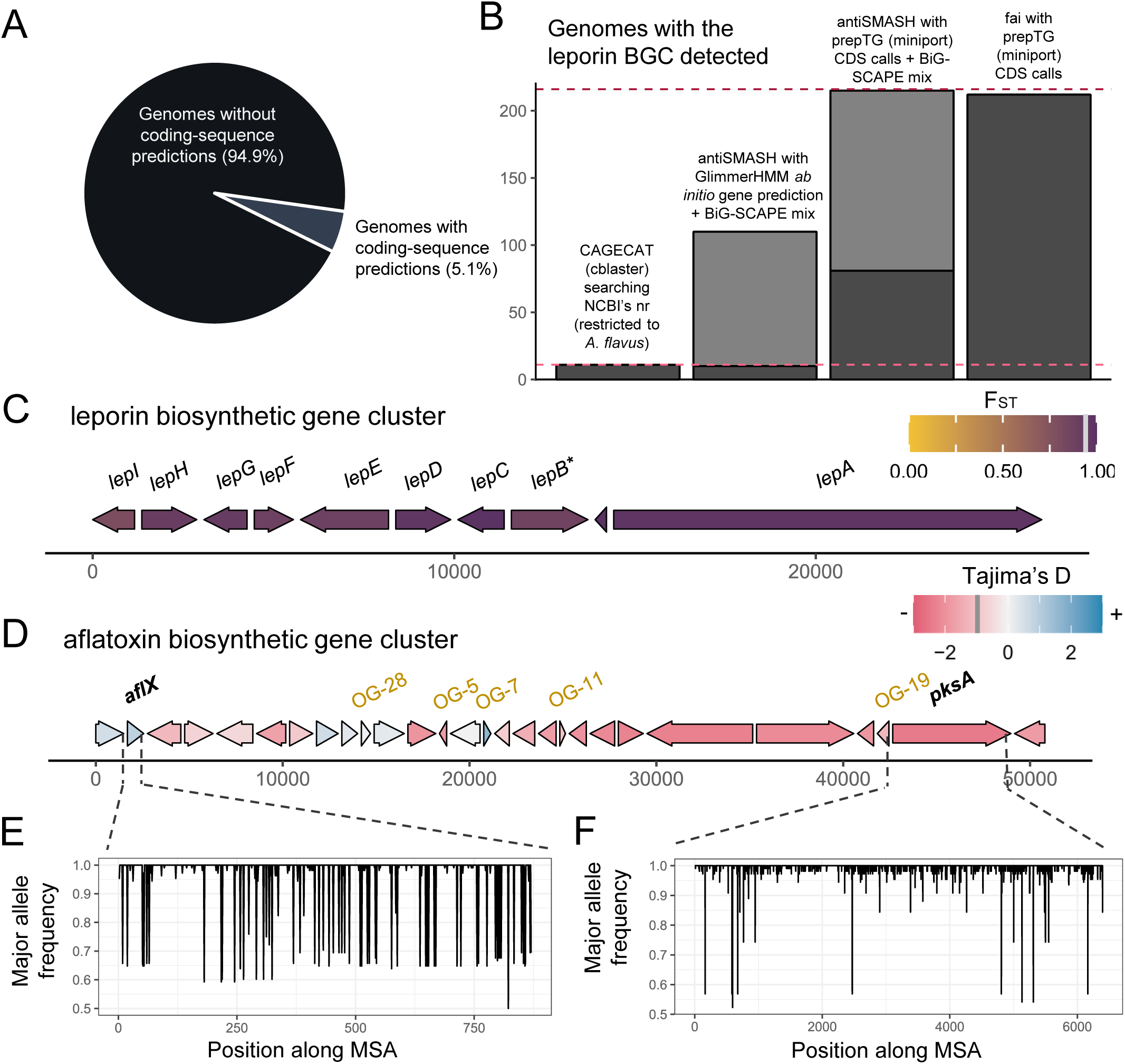
Evolutionary trends of common BGCs in *A. flavus*. **A)** The proportion of 216 A. flavus genomes from NCBI’s GenBank database with coding-sequence predictions available. **B)** Comparison of the sensitivity of fai and alternate approaches based on assemblies for detecting the leporin BGC. The dashed violet line indicates the total number of genomes (n=216) assessed and the dashed pink line indicates the number of genomes with CDS features available on NCBI (n=11). Dark grey indicates instances identified by CAGECAT/cblaster or fai or as belonging to the same GCF as the reference leporin BGC from MIBiG by antiSMASH and BiG-SCAPE analysis. Lighter grey indicates the number of similar BGCs identified by BiG-SCAPE, belonging to the same clan but not to the same GCF as the reference leporin BGC. A schematic of the (**C**) leporin and (**D**) aflatoxin BGCs is shown with genes present in ≥ 10% of samples shown in consensus order and relative directionality. Coloring of genes in (**C**) corresponds to FST values and in (**D**) to Tajima’s D values, as calculated by zol. Grey bars in the legends, at (C) 0.92 and (D) −0.98, indicate the mean values for the statistics across genes in the BGC. *For the leporin BGC, *lepB* corresponds to an updated open-reading frame (ORF) prediction by Skerker *et al.* 2021 which was the combination of AFLA_066860 and AFLA_066870 ORFs in the MIBiG entry BGC0001445 used as the query for fai. For the aflatoxin BGC, ORFs which were not represented in the MIBiG entry BGC0000008 but predicted to be within the aflatoxin BGC by mapping of gene-calls from *A. flavus* NRRL 3357 by Skerker et al. 2021 are shown in gold. The major allele frequency distributions are shown for (**E**) *aflX* and (**F**) *pksA*, which depict opposite trends in sequence conservation according to their respective Tajima’s D calculations.

Of the 212 genomes with the leporin BGC identified by fai, 202 contained instances that were high-quality and not near scaffold edges. This set of 202 instances of the gene cluster was further investigated using zol with options to perform comparative investigation of BGC instances from *A. flavus* population B genomes to instances from other populations. High sequence conservation was observed for all genes in the leporin gene cluster as previously reported^37^ (**Table S3**). Further, alleles for genes in the BGC from population B genomes were generally more similar to each other than to alleles from outside the population, as indicated by high F_ST_ values (>0.85 for 9 of 10 genes) (**Figure 3C; Table S3**). While regulation of secondary metabolites in *Aspergillus* is complex^89^, zol analysis showed that the three essential genes for leporin production^85^ also had the lowest variation in the 100 bps upstream their exonic coordinates (**Figure S5**). This suggests higher variability is occurring in the transcription of the accessory *lep* genes within the species. This supports experimental evidence that has shown gene knockouts depleting certain leporin species will still permit the production of others^85^.

fai and zol were also applied to the BGC encoding aflatoxin across *A. flavus*^90^ (Table S4). Similar to the leporin BGC, the aflatoxin BGC was highly prevalent in the species and found in 71.8% of genomes. However, in contrast to the leporin BGC, the aflatoxin BGC contained several genes with positive Tajima’s D values, indicating greater sequence variability for these coding regions across the species (**Figure 3D**). One of the genes with a positive Tajima’s D value was *aflX*, which has been shown to influence conversion of the precursor veriscolorin A to downstream intermediates in the aflatoxin biosynthesis pathway^91^ (**Figure 3E**). An abundance of sites with mid-frequency alleles in the oxidoreductase encoding gene could represent granular control for the amount of aflatoxin relative to intermediates produced. The polyketide synthase gene *pksA* had the lowest Tajima’s D value of −2.4, which suggests it is either highly conserved or under purifying selection (**Figure 3F**). In addition, because the reference proteome used to infer genomic coding regions was constructed recently^86^, fai and zol detected several highly conserved genes within the aflatoxin BGC that are not represented in the original reference gene cluster input for fai^52^. This includes a gene annotated as a noranthrone monooxygenase and recently characterized as contributing to aflatoxin biosynthesis^92,93^ (**Figure 3D**).

### Identification of the Enterococcal polysaccharide antigen and assessment of context restricted orthology inference

To demonstrate the ability of zol and fai to reliably identify ortholog groups across multiple species and thousands of genomes, we used the tools to assess the distribution of the enterococcal polysaccharide antigen (Epa) and its individual genes across the diverse genus of *Enterococcus*. Because previous comparative genomic investigations have been performed between *epa* loci from different species^94,95^, we also showcase how such prior insight can be used to tailor parameters in fai for searching for the locus across the full genus and how results from fai can be assessed for appropriate selection of parameter values in zol.

The Epa is a signature component of the cellular envelope of multiple species within *Enterococcus*^94–97^ and has mostly been characterized in the species *Enterococcus faecalis*^96,98–101^. While molecular studies have provided evidence that the locus contributes to enterococcal host colonization^100^, evasion of immune systems^102^, and sensitivity to antibiotics^103^ and phages^103,104^, it was only recently that the structure of Epa was resolved and a model for its biosynthesis and localization formally proposed^101^. A homologous instance of the *epa* locus was identified in the other prominent pathogenic species from the genus, Enterococcus *faecium*^94,95,105^; however, the prevalence and conservation of *epa* across the diverse genus of *Enterococcus*^106–108^ remains poorly studied.

We first assessed the performance of fai and zol to identify *epa* loci across representative genomes for each of the 92 species of *Enterococcus* in GTDB R214^57^ and subsequently delineate protein ortholog groups relative to other methods. Specifically, we compared the runtime and ortholog group predictions of fai and zol to the combination of cblaster and clinker as well as OrthoFinder, an established software for multi-species ortholog group delineation, run on full genomes. For this comparison, the parameter settings for fai and cblaster as well as zol and clinker were adapted to match each other more closely, with an exception being to run fai in draft-mode, which lacks an analogous feature in cblaster. The combination of fai and zol was the fastest of the three methods tested and able to identify ortholog groups for the *epa* locus in approximately one minute (**Figure 4A, S6**). Orthology inferences from fai and zol exhibited high overlap with orthology predictions by the alternate two methods, finding 96.3% of ortholog protein pairs identified by at least two of the three methods (**Figure 4B**). We also applied all three methods to determine *epa* locus orthologs across low quality representative genomes for each species to demonstrate the convenience of fai’s ability to be run in “draft mode” and improve sensitivity for detecting fragmented gene clusters in comparison to cblaster. fai identified 2.1-fold more exclusive ortholog pairs in common with OrthoFinder, expected to be relatively robust to the effects of assembly fragmentation, than the number of ortholog pairs shared exclusively by cblaster and clinker with OrthoFinder (**Figure 4C**). In addition, we performed evolutionary-simulation of the *epa* locus, allowing for sequence gains and losses, and assessed context-limited orthology inference by zol, clinker and OrthoFinder (**Figure S7; Supplementary Text**). zol was able to recover a high fraction of true positive ortholog relations and was the best method at avoiding prediction of false positive orthologs.

**Figure 4:**
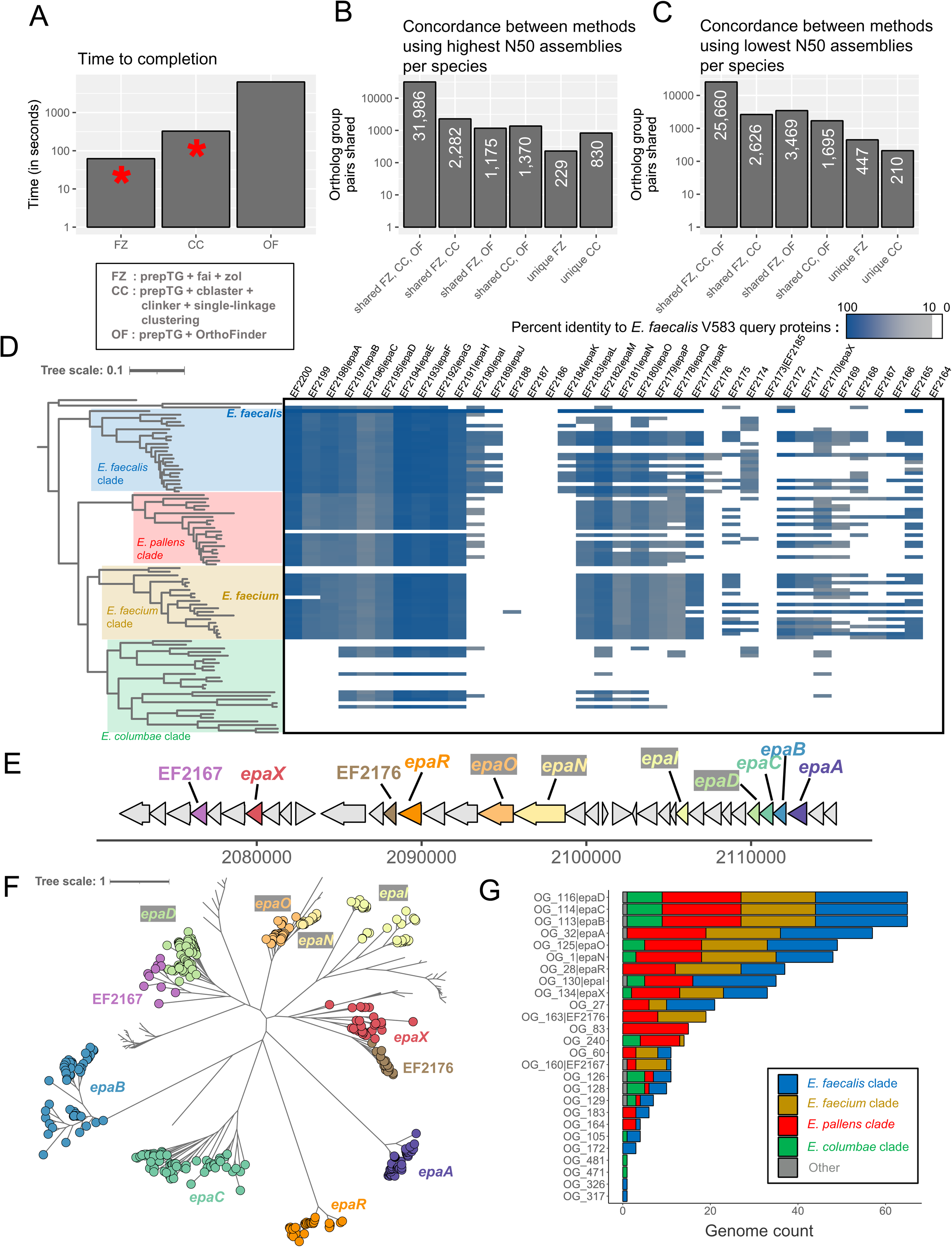
Searching for the *epa* locus across the diverse genus of *Enterococcus*. **A**) Overview of the time needed to run orthology/homology inference methods on the 92 genomes with the highest N50 for each distinct *Enterococcus* species. OrthoFinder was run at the genome-wide scale, while fai and cblaster were used to first identify genomic regions corresponding to the *epa* locus from *E. faecalis* V583 and subsequently zol and clinker were applied to determine ortholog groups, respectively. The red asterisks denote that manual assessment or filtering of homologous gene clusters identified by fai and cblaster is encouraged and thus additional time might be required for them. Counts showing the overlap in orthologous protein pair predictions by the three different methods are shown following their application to representative genomes from GTDB R214 with the **B**) highest N50 and **C**) lowest N50 for the 92 different species. **D**) The distribution of the *epa* locus, based on criteria used for running fai, is shown across a species phylogeny for 92 genomes representative of distinct *Enterococcus* species in GTDB R214. The coloring of the heatmap corresponds to the percent identity of the best matching protein from each genome to the query *epa* proteins from *E. faecalis* V583. **E**) A schematic of the *epa* gene cluster from *E. faecalis* V583 (from EF2164 to EF2200) with glycosyltransferase encoding genes shown in color. **F**) A maximum-likelihood phylogeny of zol-identified ortholog groups corresponding to glycosyltransferases in *epa* loci across *Enterococcus*. **G**) Distribution of different glycosyltransferase ortholog groups across the four major clades of *Enterococcus* are shown. For **D** and **F** the tree scales correspond to the number of amino acid substitutions along the alignments used for phylogeny construction.

Next, to properly and comprehensively assess the distribution of *epa* across the entire set of 5,291 genomes in GTDB classified as one of the 92 *Enterococcus* species^57^, we applied fai with more careful consideration of parameter values and requested more advanced features for gene cluster detection. A sensitive searching criterium was selected based on prior comparative genomics for the locus^94,95^ and its coordinates along the *E. faecalis* V583 genome as a reference^99,101^. For detection of *epa* orthologous regions, co-location of at least seven of the 14 *epa* genes previously identified as conserved in both *E. faecalis* and *E. faecium* was required. The default threshold for syntenic conservation of homologous instances to the query gene cluster was disregarded to increase sensitivity for the detection of *epa* in enterococcal species more distantly related to *E. faecalis*. In addition, key proteins were specified and the length of the flanking context to include as part of the loci was expanded. Using these criteria, 5,085 of the genomes assessed were found to possess an *epa* locus, with phylogenomic investigations further revealing that the locus is highly conserved in three of the four major clades of *Enterococcus* (**Figure 4D; Table S5**).

Based on fai’s reports, we realized that to achieve optimal clustering for ortholog groups across the diverse set of *epa* loci identified, we needed to lower the default thresholds for percent identity and coverage that protein pairs needed to exhibit for being considered as orthologs (**Figure 4D; Table S5**). We ran zol on both the full set of 5,052 high-quality *epa* loci and only loci from species representative genomes. For the comprehensive analysis, zol was able to identify 14 ortholog groups as core or near-core, found in >90% of loci instances (**Table S6**). When provided 30 threads, zol completed in 30.7 hours and had a maximum memory usage of 101.3 GB. The more restricted analysis of zol to investigate *epa* instances from 65 species representative genomes was to allow for assessing the quality of ortholog group predictions using phylogenetics (**Table S7**). After applying zol on *epa* from species representative genomes, orthology predictions were assessed through construction of a maximum-likelihood phylogeny of *epa* associated glycosyltransferases. Ortholog groups which corresponded to glycosyltransferases from *E. faecalis* V583 were labelled on the phylogeny and confirmed to match distinct phylogenetic clades, which suggests their appropriate delineation (**Figure 4EF**). zol further identified several *epa* associated glycosyltransferase ortholog groups that were absent in the *E. faecalis* representative genome and other representative genomes from the *E. faecalis* clade (**Figure 4G**). These distinct glycosyltransferases might impact the final structure or decoration of Epa in other *Enterococcus* species.

### zol identifies genetic diversity of *epa*X-like glycosyltransferases in *E. faecalis*

zol features several options related to the dereplication of input gene clusters to retain only distinct representative instances for orthology inference and other downstream analytics (**Figure S8**). Importantly, the application of these methods can substantially reduce zol’s runtime and impact some of the evolutionary statistics computed (**Figure S8, S9, S10, Supplementary Text**). Whether dereplication is appropriate for a particular analysis should thus be carefully considered by users depending on their research aims. In particular, dereplication can impact investigations for highly sequenced bacterial taxa, including the opportunistic pathogen *E. faecalis*. For such pathogens, certain lineages, such as those commonly isolated at clinics, might be overrepresented in genomic databases, and the researcher may find it beneficial for the analysis to apply dereplication.

To showcase the scalability of zol and its ability to expand knowledge for even well-studied gene clusters, we applied it to high-quality, complete *epa* loci from 1,232 *E. faecalis* genomes without dereplication. In accordance with prior studies^94,101^, zol was able to distinguish core and strain-variable patterns. The report from zol showed that one end of the locus corresponds to genes which are highly conserved and core to *E. faecalis* (*epa*A-*epa*R), whereas the other end contained strain-specific genes (**Figure 5A; Table S8**). Using zol, we further found that variably conserved genes exhibit high sequence dissimilarity, as measured using both Tajima’s D and average sequence entropy, in comparison to the core genes of the locus (**Figure 5BC**). These statistics were robust to the application of dereplication and thus unlikely to be heavily impacted by well-sequenced lineages (**Figure S9, S10**).

**Figure 5:**
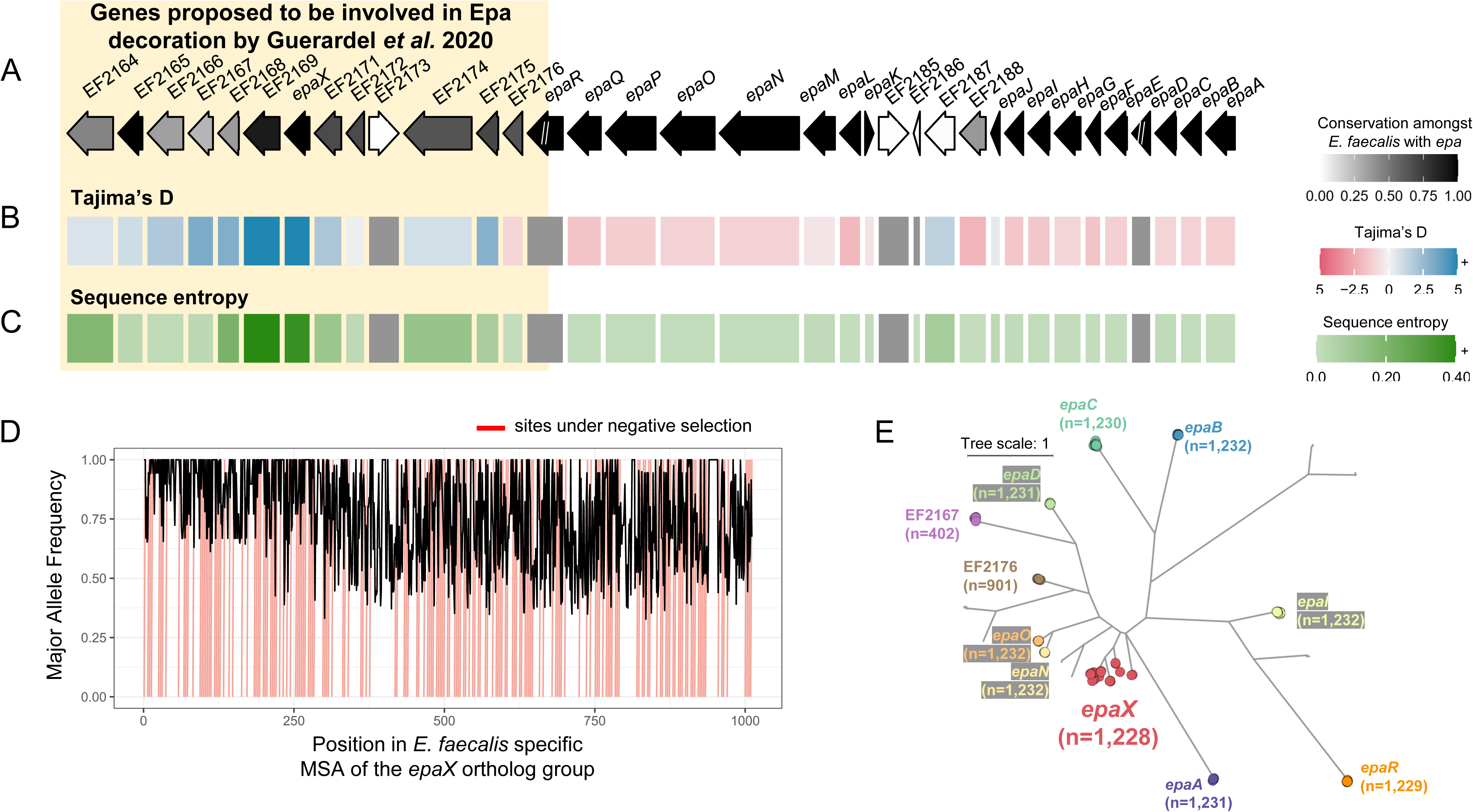
High sequence diversity of *epaX*-like glycosyltransferases amongst *E. faecalis*. A schematic of the *epa* locus from *E. faecalis* V583 with evolutionary statistics, **A**) conservation, **B**) Tajima’s D and **C**) sequence entropy, gathered from the best corresponding ortholog group for each protein. Ortholog groups were inferred from zol investigation of 1,232 *epa* loci from the species. Genes upstream of and including *epaR* were recently proposed to be involved in Epa decoration by Guerardel *et al.* 2020. “//” indicates that the ortholog group was not single-copy in the context of the gene-cluster and calculation of evolutionary statistics for these genes was avoided (grey in panels B and C). Note, the same ortholog group was regarded for EF2173 and EF2185 which correspond to an identical *ISEf1* transposase. The length of proteins in the locus schematic are the median lengths of the corresponding ortholog groups. **D**) The major allele frequency is depicted across the alignment for the ortholog group featuring *epaX*. Sites predicted to be under negative selection by FUBAR, Prob(*α*>*β*) ≥ 0.9, are marked in red. **E**) An approximate maximum-likelihood phylogeny of glycosyltransferase ortholog groups identified by zol which were found in >1% of *epa* instances. Ortholog groups identified by zol are indicated by colored circular nodes with names of *epa* genes from *E. faecalis* V583 noted where possible. The number of leaves/proteins for each clade is provided for labeled ortholog groups. The tree scale corresponds to the number of amino acid substitutions along the input protein alignment used for phylogeny construction.

One ortholog group, corresponding to the glycosyltransferase *epaX*, exhibited substantially higher sequence variation than other *epa* associated glycosyltransferases (**Figure 5BD**). This finding was further validated through phylogenetic analysis of glycosyltransferases from the species, which highlighted the breadth of diversity observed for the *epaX* ortholog group relative to other *epa* associated glycosyltransferases (**Figure 5E**).

## Discussion

Here fai and zol are introduced to enable large-scale evolutionary investigations of gene clusters in diverse taxa. Together these tools overcome current bottlenecks in computational biology to infer orthologous sets of genes at scale across thousands of diverse genomes and large metagenomic assemblies.

The set of input gene clusters for zol does not need to be produced by fai. cblaster^45^ is another tool that can identify instances of a query gene cluster within a set of target genomes and extract them in GenBank format for downstream investigations using zol. For those lacking computational resources needed for fai analysis, cblaster offers remote searching of BGCs using NCBI’s BLAST infrastructure and non-redundant databases. More recently, CAGECAT^87^, a highly accessible web-application for running cblaster, was also developed and can similarly be used to identify and extract gene cluster instances from genomes represented in NCBI databases. In contrast to these tools, prepTG and fai feature algorithms and options for users interested in: (i) identification of gene clusters in metagenomes, (ii) performing standardized gene annotation across target genomes, (iii) improved sensitivity for gene cluster detection in draft-quality assemblies, and (iv) automated filtering of secondary, or paralogous, matches to query gene clusters. In addition, users can apply zol to further investigate homologous sets of gene clusters identified from IslandCompare^109^, BiG-SCAPE^44^, or vConTACT2^110^ analyses, which perform comprehensive clustering of predicted genomic islands, BGCs, or viruses.

The application of fai to identify gene clusters in metagenomes is demonstrated here through rapid, targeted detection of a virus across lake metagenomic assemblies. We expect that both fai and zol will gain greater relevance for metagenomic applications in the future as long-read sequencing becomes cheaper. Importantly, the tools can be applied directly on assemblies without the need for binning scaffolds into MAGs, avoiding complications associated with binning^111^. In addition to their application to viral tracking, fai and zol’s application to metagenomes could be useful for assessing the presence of concerning transposons carrying antimicrobial resistance traits^112–114^ and identifying novel auxiliary genes within known BGCs which may tailor the resulting specialized metabolites and expand chemical diversity^115,116^.

Reidentifying gene clusters in eukaryotic genomes remains difficult due to technical challenges in gene prediction owing to the presence of alternative splicing. The ability of fai and zol to perform population-level genetics on BGCs from the eukaryotic species *A. flavus* was demonstrated. While there are over 200 genomes of *A. flavus* in NCBI, only 5.1% have coding-sequence information readily available. We used miniprot^55^ to map high quality gene coordinate predictions from a representative genome in the species^86^ to the remainder of genomic assemblies with prepTG which enabled high sensitivity detection of BGCs with fai. Our analysis provides additional support that the leporin BGC is conserved across the species^37^ using an assembly-based approach.

The ability of zol to identify ortholog groups across 5,052 gene cluster instances from 71 distinct species using limited computational resources was demonstrated through investigation of the *epa* locus across *Enterococcus*. While such large-scale investigations will be largely limited to those with access to a server, we expect datasets to often feature some degree of species level redundancy. For instance, 80.2% of the 5,052 *epa* instances were from only two species, *E. faecalis* and *E. faecium*. Thus, to alleviate computational costs, we have included functions for dereplication of gene clusters and reinflation of ortholog groups in zol. Applying these features to the comprehensive set of *epa* loci using 30 threads, reduced runtime from 30.7 to 3.5 hours and maximum memory usage from 101.3 GB to 83.2 GB (**Table S9**).

We further assessed the quality of ortholog group predictions by fai and zol using phylogenetic investigations and comparisons with other software for homology inference. Specifically, we compared orthology inference results from fai and zol to predictions obtained from the combination of cblaster and clinker as well as OrthoFinder^117^, which was used to detect ortholog groups at the genome-wide scale. Notably, clinker^46^, which is developed by the authors of cblaster, is primarily designed to produce interactive visualizations showing relationships between related gene cluster instances. clinker’s application of single-linkage clustering to determine related sets of genes and to color matching genes in figures is expected to produce relatively coarse ortholog groups. OrthoFinder was chosen as a representative method for standard multi-species orthology inference because it has been shown to perform well for several criteria in prior benchmarking studies^117,118^. Through application to identification of ortholog groups for diverse *epa* loci from multiple distinct species and evolutionary simulation of the locus from *E. faecalis*, we found zol produces reliable orthology predictions that are mostly in accordance with alternate orthology inference methods while exhibiting restraint for over clustering. In the future, we are considering further improving the algorithm for ortholog group classifications within zol. Specifically, we might take a similar approach to OrthoFinder in which coarse ortholog groups are first identified and later refined using phylogenetics.

Our investigation of *epa* loci from multiple species revealed the presence of a multitude of glycosyltransferases associated with production or decoration of the polysaccharide, including some that are absent in the representative *E. faecalis* genome, the species in which the polysaccharide has been most extensively characterized. Through population-genetic investigations of the locus in *E. faecalis* using zol, we further determined that an ortholog group containing *epa*X-like glycosyltransferases possessed high sequence divergence relative to other glycosyltransferases associated with the locus. In addition to influencing the ability of *E. faecalis* to colonize hosts^100^, mutations in *epa*X and other genes from the ortholog group have also been shown to impact susceptibility to phage predation^119–122^. Therefore, we hypothesize that extensive evolution of the *epa*X ortholog group is a result of contrasting selective forces, pressuring *E. faecalis* to retain or (re-)acquire the glycosyltransferase to gain a fitness advantage within hosts but also lose the gene to escape phage predation.

## Conclusions

Practically, zol presents a comprehensive analysis tool for comparative genetics of related gene clusters to facilitate detection of evolutionary patterns that might be less apparent from visual analysis. Fundamentally, the algorithms presented within fai and zol enable the reliable detection of orthologous gene clusters, and subsequently orthologous proteins, across multi-species datasets spanning thousands of genomes and help overcome a key barrier in scalability for comparative genomics.

## Methods

### Software availability

zol is provided as an open-source software suite, developed primarily in Python3 on GitHub at: https://github.com/Kalan-Lab/zol. Docker and Bioconda^123^ based installations of the suite are supported. For the analyses presented in this manuscript, we used v1.4.1 of the zol software package^124^. Version information for major dependencies of the zol suite^53,55,62,65,125–132^ and other software used^44,74,133^ for analyses in this study is provided in Table S10. Code and input files for generation of figures in this manuscript are provided separately on GitHub at: https://github.com/Kalan-Lab/Salamzade_etal_zol.

### Availability of data and materials

Genomes and metagenomes used to showcase the application of fai and zol are listed with GenBank accession identifiers in Table S11. Total metagenomes and their associated information from Lake Mendota microbiome samplings were originally described in Tran *et al.* 2023^73^ and deposited in NCBI under BioProject PRJNA758276. Genomic assemblies available for *A. flavus* in NCBI’s GenBank database on Jan 31st, 2023 were downloaded in FASTA format using ncbi-genome-download (https://github.com/kblin/ncbi-genome-download). Genomic assemblies for *Enterococcus* that met quality and taxonomic criteria for belonging to the genus or related genera (e.g. Enterococcus_A, Enterococcus_B, etc.) in GTDB^57^ release R207 were similarly downloaded from NCBI’s GenBank database using ncbi-genome-download in FASTA format.

#### Assessment of compute time, memory usage, and disk space

The UNIX *time* command was applied to measure the runtime and memory usage of programs. Specifically, the “Elapsed (wall clock) time” was regarded as the runtime and the “Maximum resident set size (kbytes)” as the maximum memory usage. The UNIX *du* command was used to measure the final disk space used by various programs. All analyses were computed on the same server running Ubuntu 18.04.06 LTS with AMD EPYC 7451 24-Core processors, 472 GB of 288-Pin DDR4 random-access memory, and a Samsung 970 Pro solid disk drive.

### Overview of tools and algorithms

#### prepTG - processing and preparing target genomes for searching with fai

prepTG allows users to create a database of target genomes that can be searched for homologous instances of query gene clusters with fai. In addition to formatting and producing files for optimizing fai searches, prepTG integrates pyrodigal^53^, prodigal^54^, and miniprot^55^ for gene-calling or protein-mapping in prokaryotic and eukaryotic genomes as well as metagenomes to aid consistency in fai’s performance and limit bias due to potential differences in gene-calling methods. For miniprot-based protein-mapping, coding sequence predictions are required to exhibit an identity of at least 80% to the reference protein and instances of overlapping mRNA and exon features are resolved by retaining only the highest scoring mappings.

prepTG also features options to download pre-built databases for select bacterial taxa that are commonly studied^56^, such as ESKAPE pathogens, or to download all genomes belonging to any genus or species in GTDB R214^57^ and subsequently construct a database *ab initio*.

#### fai - automated identification of homologous instances of gene clusters

fai allows for rapid detection of gene clusters in target genomes. It accepts a target genomes database prepared by prepTG and query gene cluster(s). Query gene cluster(s) can be provided in one of three formats: (i) GenBank file(s) with CDS features, (ii) a coordinate along a reference genome, or (iii) a set of proteins. When using coordinates along a reference genome to define a gene cluster, fai reperforms gene-calling along the reference using pyrodigal^53^ and extracts a local GenBank file corresponding to the specified region.

zol implements HMM-based and CDS separation-based approaches for determining homologous gene cluster instances in target genomes, which can further be combined in a hybrid approach. For both approaches, homologs of proteins from query gene clusters are first searched for in predicted proteomes of target genomes using DIAMOND alignment^130^. Then, in “Gene-Clumper” mode, which is the default, scaffolds with homologs of query proteins are dynamically assessed for whether homologs are within a maximum number of CDS predictions to be regarded as belonging to the same gene cluster. In “HMM” mode, scaffolds of target genomes are instead scanned gene-by-gene using an HMM and neighborhoods or sets of genes are regarded as being in a state of homology to the query gene cluster if several individual genes depict homology to the proteins from the query gene cluster(s). The algorithm is similar to *lsa*BGC-Expansion^38^, however, it is not dependent on a preliminary genome-wide orthology grouping analysis and thus features a different set of filters to still enable high-throughput automated detection of homologous gene cluster segments as a result. *lsa*BGC-Expansion is reliant on a preliminary orthology analysis to identify BGC-specific genes that could be used to differentiate true homologous instances of BGCs and customize weighting of HMM emission probabilities for distinct genes. It further requires the length of genes within putative homologous regions to be within a certain deviation from the median length of known gene instances. In contrast, fai has preconfigured emission probabilities which can be customized by users and has no length requirement for potential homologous instances of genes. fai further allows the “HMM-based” approach to be run with the parameter for aggregating CDS predictions for the “Gene-Clumper” mode, whereby, gene cluster segments detected by the HMM can be joined with other such segments if they are withing a certain number of CDS features from each other. Similar to *lsa*BGC-Expansion, syntenic similarity between candidate and query gene cluster segments can also be used to filter candidate segments using a gene cluster-wide correlation metric^38^.

By default, fai requires filters pertaining to the number of genes from query gene clusters to be met for each homologous gene cluster candidate segment. However, in “draft mode”, thresholds for detection of gene clusters within target genomes are assessed in aggregate for putative gene cluster segments found near scaffold edges (< 2,000 bp). Visual reports produced by fai showcasing the sequence similarity of target genome proteins to the query protein(s) can then be manually investigated by users to assess the validity of fragmented gene cluster instances. In addition, fai features an option to filter for paralogous, overlapping candidate segments of a gene cluster in target genomes and offers an intuitive visualization of gene cluster segments, if requested, to allow users to assess their quality, including proximity of candidate segments to scaffold edges. Together, these options enable the large-scale identification of orthologous gene clusters across genomes which can then be leveraged by zol to perform context-specific inference of protein ortholog groups.

In addition to a directory of homologous gene clusters in GenBank format, to serve as input for zol analysis, and a small set of visual PDF files, fai generates an in-depth report on which target genomes have the query gene cluster as an XLSX spreadsheet. This spreadsheet includes information such as the average amino acid identity (AAI), syntenic similarity, and number of conserved genes for gene clusters from target genomes relative to the query gene cluster. The spreadsheet allows for easy sorting of various columns to assist identification of which target genomes feature a gene cluster to the desired degree of similarity for the user.

#### zol - computes a variety of evolutionary statistics and can perform gene cluster specific dereplication

The zol workflow begins by processing the input directory of gene cluster GenBank files to assess validity and perform filtering of gene clusters or individual proteins. Filtering can be performed at the gene cluster level by requesting filtering of draft-quality gene clusters, those marked as being near scaffold edges, or low-quality gene clusters, those with ≥10% missing base-pairs (e.g. Ns) in their sequence. Filtering of individual proteins which are near scaffold edges can also be performed if fai was used to identify the input gene cluster set, because fai marks these proteins with a special feature tag in the resulting gene cluster GenBank files.

Next, zol will perform dereplication of gene clusters, if requested by users, with skani^65^ by clustering gene clusters which depict some user-defined coverage and identity thresholds using single linkage clustering or more resolved MCL-based clustering, for which the inflation parameter can be adjusted. Representative gene clusters are selected from each cluster as part of the dereplication based on maximum length and, if comparative analysis is requested, whether the representative gene cluster is part of the focal or focal-complement set of gene cluster instances specified by the user.

The input set of gene clusters or set of dereplicated representative gene clusters is then used to identify protein ortholog groups with an InParanoid-type approach^3^. Briefly, DIAMOND^130^ is used to perform all vs. all pairwise alignment between proteins from the set of gene clusters after which the alignments are processed to identify reciprocal best hits (RBH) between pairs of gene clusters. In-paralogs are identified within each gene cluster based on whether two coding sequences depict more similarity to each other than one does to an RBH with a different gene cluster. Bitscores, standardized through division by reflexive bitscore values for query proteins, are used to assess homology. Specifically, the average normalized bitscore between each pair of orthologs and in-paralogs is recorded. Afterwards, bitscores between such protein pairs are further standardized through dividing them with the average values between pairs of gene clusters to aid proper clustering of proteins downstream. This is akin to the genome-wide normalization procedure recommended in OrthoMCL, owing to the realization that orthologs between distantly related species are also more likely to exhibit lower sequence similarity, which should be corrected for prior to MCL clustering^2^. This information is input into MCL with the inflation parameter set to 1.5, similar to other orthology inference methods^7,117^. The inflation parameter and minimum identity and coverage cutoffs to consider valid pairs of in-paralogs and orthologs are adjustable by users.

Reinflation can also be requested by users to expand ortholog groups to include proteins from the full input set of gene clusters if gene cluster dereplication was requested^10^. Reinflation of ortholog groups is performed by first performing comprehensive and granular clustering of proteins from all input gene clusters using CD-HIT^128^, requiring proteins to depict >98% sequence similarity and > 95% bi-directional coverage to the representative sequences of clusters. Proteins in CD-HIT clusters are then mapped to ortholog groups if they co-cluster with proteins from dereplicated gene clusters which are already assigned to ortholog groups. Dereplication and reinflation are not recommended if sequence redundancy amongst the set of input gene clusters is low. Stringent cutoffs used for CD-HIT clustering during reinflation assume that dereplication was also run with stringent parameters to only collapse highly similar gene clusters. Otherwise, reinflation could miss more distant instances of ortholog groups, resulting in an underestimation of ortholog group conservation amongst gene clusters.

Next, zol will partition protein and nucleotide sequences from gene clusters according to ortholog groups, perform protein alignment using MUSCLE^132^, and create codon alignments using PAL2NAL^134^. We also offer an option to use reference proteins to refine and filter sequences based on multiple sequence alignment using MUSCLE^132^, which might be useful to further filter intronic sequences in eukaryotic ORFs. Codon alignments are filtered for regions with high ambiguity (≥10% gaps) using trimAL^126^ which are then used downstream for calculation of evolutionary statistics and to construct approximate maximum-likelihood phylogenies using FastTree 2^127^ for each ortholog group. Consensus protein sequences for each ortholog group are finally constructed using HMMER3^129^.

Using protein consensus sequences of each ortholog group, zol is next able to linearize annotation of ortholog groups with various annotation databases including KOfam^14^, the PGAP database^135^, VFDB^51^, CARD^61^, MIBiG^52^, ISfinder^60^, the PaperBLAST database^136^, and Pfam^137^. A custom FASTA file can also be provided by users to annotate ortholog groups. The best hit per ortholog group for each annotation database is selected by score, if annotation is HMM based^138^, or bitscore, if it is DIAMOND alignment based^130^, and a default E-value cutoff of 1e-5. The E-value of the alignment is provided in the zol report for each putative annotation except Pfam domains. However, for Pfam annotations, only domains meeting trusted thresholds are reported.

Next, zol will compute basic statistics per ortholog group including the consensus order, consensus directionality, whether proteins are single-copy across gene clusters, the median length of ortholog group sequences, their median GC% percentage, and GC skew values. The consensus order and directionality are performed similarly to *lsa*BGC-PopGene^38^. Afterwards, in the sixth step, zol will calculate evolutionary statistics for each ortholog group including Tajima’s D^49^, the proportion of filtered codon alignments which correspond to segregating sites, the average sequence entropy of the filtered codon alignment and the 100 upstream region, and the median and maximum Beta-RDgc. Beta-RDgc is a statistic that is derived from the Beta-RD statistic which we described in *lsa*BGC^38^ and measures the divergence of a pair of protein sequences based on the expected divergence between the gene clusters. Values below one suggest that protein divergence is larger for the pair than expected based on other shared proteins between the two gene clusters; conversely, the opposite trend might suggest high conservation of the particular protein between the gene clusters and potentially gene-specific horizontal gene transfer. Finally, we perform site-specific selection analyses using the FUBAR^139^ and GARD^140^ methods offered in the HyPhy suite. While highly scalable relative to comparable methods^139^, these analyses can still take considerable time and are turned off by default. Importantly, GARD recombination detection^140^ and partitioning of input alignments for ortholog groups can also be used for alternate HyPhy analyses with HyPhy Vision^62^, to extend beyond the site-specific selection analyses using FUBAR^139^ supported directly in zol.

Prior to the generation of a final report, zol allows users to perform an optional comparative analysis between user-defined set(s) of focal and complementary or alternate gene cluster instances. In these comparative analyses, the conservation and fixation index^70^ is calculated for each ortholog group.

Finally, we generate a consensus report and a spreadsheet in XLSX format where each row corresponds to an ortholog group and columns correspond to basic statistics, evolutionary statistics, and annotation information. Quantitative fields are automatically colored to make visual detection of patterns easier for users. A basic heatmap showing the presence of ortholog groups across gene clusters is also produced.

zol additionally features two alternate modes that can be triggered via specific arguments. First, the “only-orthologs” argument will invoke zol to only compute ortholog groups and exist after determining them. Second, the “select_fai_params_mode” argument allows users to provide a handful of known instances for a gene cluster and determine appropriate thresholds for searching for additional instances of the gene cluster using fai. This mode assumes that the known instances provided are representative of the breadth of diversity expected for the gene cluster amongst the target genomes being searched.

#### abon, atpoc, and apos – tools for assessing novelty and conservation of BGCs, phages, and plasmids from a single strain

The zol suite features three small wrapper programs called abon, atpoc, and apos which assess the conservation and novelty of a single genome’s BGC-ome, phage-ome, and plasmid-ome, respectively, relative to a target genome database constructed by prepTG. The target genomes database could be all other genomes belonging to the focal genome’s species or genus. The three programs are wrappers of fai but also offer a simple BLAST search alternative, to more thoroughly check for whether individual genes from BGCs, phages, and plasmids are present in the target genomes being searched. These tools accept results from standard software for annotation of BGCs^133,141^, phages^74,142,143^, and plasmids^143,144^ but do not integrate them within the suite. Similar to fai and zol they produce auto-formatted XLSX spreadsheets as primary results.

### Application of fai and zol to track a virus within lake metagenomes

VIBRANT was used to identify viral contigs or sub-contigs in the three total metagenomes from Tran *et al.* 2023^73^ sampled on the earliest date of 07/24. Afterwards, predicted circular contigs were clustered using BiG-SCAPE^44^ which revealed a ∼36 kb virus was found in two of the three metagenomes.

prepTG was run on all 16 total metagenomic assemblies from the Tran *et al.* 2023 study, performing gene calling with pyrodigal in metagenomics mode^53^ to prepare for comprehensive targeted searching of the virus with fai. fai was run with largely default settings, with filtering of secondary instances of the virus requested to retain only the best matching scaffold or scaffold segment resembling the queries. In addition, the syntenic correlation requirement of hits to the query gene clusters was turned off to account for the circular nature of the virus, which the assessment is not designed for. To assess the performance of cblaster for preparing the target metagenomes database and subsequently searching for the virus, we provided GenBank files with CDS features produced by prepTG as input for cblaster makedb and adjusted searching parameters for cblaster search to more closely match what we used for fai.

### Microevolutionary investigations of leporin and aflatoxin BGCs in *Aspergillus flavus*

Genomic assemblies downloaded from NCBI GenBank were processed using prepTG. Of the 217 genomic assemblies downloaded, one, GCA_000006275.3, was dropped from the analysis because the original GenBank file had multiple CDS features with the same name, leading to difficulties in performing BGC prediction with antiSMASH^133^, and because alternate assemblies were available for the isolate. prepTG was run on all assemblies with miniprot^55^ based gene-mapping of the high-quality gene coordinate predictions available for *A. flavus* NRRL 3357 (GCA_009017415.1)^86^ requested. Target genomes were then searched for the leporin (BGC0001445) and aflatoxin (BGC0000008) BGCs using GenBank files downloaded from MIBiGv3^52^ as queries. For leporin, AFLA_066840, as represented in the MIBiG database, was treated as a key protein required for detection of the BGC. Similarly, for aflatoxin, PksA (AAS90022.1), as represented in the MIBiG database, was treated as a key protein required for detection of the BGC. Draft-mode and filtering of paralogous segments was requested. For both analyses, ortholog groups found in fewer than 5% of gene cluster instances were disregarded.

We reidentified population B as previously delineated^37^ using k-mer based ANI estimation^145^ and neighbor-joining tree construction^146^. A discrete clade (n=81) in the tree was validated to feature all isolates previously determined as part of population B^37^ and thus regarded as such.

For comprehensive and *de novo* BGC prediction, antiSMASH was run on the 216 genomic assemblies with ‘glimmerhmm’ requested for the option ‘--genefinding-tool’. Similarly, antiSMASH was also run on full GenBank files for genomes generated by prepTG from reference proteome-mapping via miniprot. For one genome, antiSMASH was unable to process the full GenBank created by prepTG due to an error related to “inconsistent exon ordering”. BGCs from each set of genome annotations were independently clustered using BiG-SCAPE with “mix” clustering analysis and MIBiG reference BGC integration requested. The gene cluster family and clan matching the reference leporin BGC in MIBiG (BGC0001445) were regarded as the leporin BGC. For remote cblaster^45^ analysis, CAGECAT^87^ was used to search NCBI’s nr database with proteins from the leporin BGC representative (BGC0001445) provided as a query. Only 13 scaffolds, belonging to 12 assemblies (including GCA_000006275.3), were identified.

### Evolutionary investigations of the *epa* locus across *Enterococcus*

All *Enterococcus* genomes represented in GTDB R207^57^ (n=5,291) were downloaded using ncbi-genome-download^53^. The same query for *epa* was used for all analyses. Specifically, coordinates extending from 2,071,671 to 2,115,174 along the *E. faecalis* V583 chromosome, corresponding to genes EF2164 to EF2200, were used as a query for the *epa* locus in fai to identify homologous instances in target genomes^99,101^.

#### Comparing orthology/homology inferences between fai & zol, cblaster & clinker, and OrthoFinder

Representative genome assemblies were selected for each of the 92 species of *Enterococcus* in GTDB R214^57^ based on the N50 metric. One set of species representative genomes corresponded to those with the largest N50 values and the other set was comprised of genomes with the lowest N50 values. The two sets of species representative genomes were processed and investigated identically but independently. Gene calling was first performed for genomes using prepTG with pyrodigal^53^. To generate the input for OrthoFinder, proteins from prepTG’s genome-wide GenBank files were extracted in FASTA format. After, OrthoFinder was run with default settings. Phylogenetic hierarchical orthogroups inferred by OrthoFinder were used for comparisons. To perform gene cluster specific homology prediction with cblaster and clinker, we first used cblaster makedb to convert the genome-wide GenBank files from prepTG into a database that could be searched with cblaster search. cblaster search was run using the criteria: (i) DIAMOND alignment sensitivity mode set to very-sensitive, (ii) the percentage of query genes required to be present in a cluster set to 25%, (iii) 1e-10 as the maximum E-value for protein hits to be considered, (iv) 0% as the minimum coverage for protein hits to be considered, (v) 0% as the minimum identity for protein hits to be considered, (vi) the maximum flanking context for the gene cluster to gather set to 0 bp, (vii) request for intergenic proteins to be included, and (viii) a maximum of 4620 bp allowed to separate protein hits for them to be considered as part of the same gene cluster, which should approximately correspond to the aggregate length of 5 bacterial genes on average^147^. Next, cblaster extract_clusters was used to extract gene clusters found in target genomes by cblaster in GenBank format and provide them as input for clinker. clinker was run using default settings but with only an output and matrix output file requested to cut time needed to render an interactive figure, its primary intended result file. To aid appropriate comparisons in orthology prediction, fai was largely run using similar criteria as cblaster search: (i) DIAMOND alignment sensitivity mode set to very-sensitive, (ii) the percentage of query genes required to be present in a cluster set to 25%, (iii) 1e-10 as the maximum E-value for protein hits to be considered, (iv) the maximum flanking context for the gene cluster to gather set to 0 bp, (v) a maximum of 5 proteins allowed to separate hits for them to be considered as part of the same gene cluster, and (vi) syntenic similarity assessment between target gene clusters and the query gene cluster turned off. However, draft-mode was enabled in fai, which is not available in cblaster, to showcase the program’s ability to improve sensitivity for draft-quality assemblies. zol was applied with mostly default settings but with the flags “only-orthologs”, to stop after it determined ortholog groups, and “allow_edge_cds”, to allow usage of CDS features marked by fai to be near scaffold edges. All three methods were provided 20 threads wherever possible.

#### Comprehensive and tailored usages of fai and zol for finding epa in Enterococcus

Based on prior comparative analyses that had shown that gene conservation and gene order can be slightly variable between *epa* loci from *E. faecalis* and *E. faecium*^94,95^, we relaxed the syntenic similarity requirement of candidate gene cluster matches in target genomes to the query in fai from 0.6 to 0.0. In addition, we relaxed the minimum percentage of query proteins needed to report a homologous instance of the *epa* locus to 10%. Instead, we required the presence of 50% of key *epa* proteins found in both *E. faecalis* and *E. faecium*, defined as *epaABCDEFGHLMOPQR*, for the identification of valid homologous instances of the *epa* locus. The E-value cutoff to determine presence for the key *epa* proteins was lowered from 1e-20 to 1e-10 to be inclusive of shorter genes and allow for higher levels of sequence divergence across the *Enterococcus* genus. To gather auxiliary genes flanking the core *epa* region in target genomes, we further requested the inclusion of CDS features found within 20 kb of the boundary genes in detected instances of the *epa* locus within the resulting GenBank files produced by fai. A phylogenetic heatmap was constructed for the presence of the *epa* locus across a species tree using species representative genomes, selected based on largest assembly N50, where the values of the heatmap corresponded to the maximum percent identity of a query protein to their best match in target genomes. Because EF2173 and EF2185 are identical transposases, they were shown as one column in the heatmap. The species tree was constructed using GToTree^148^ using HMMs for proteins regarded as largely single-copy core to the phylum Bacillota. The phylogenetic heatmap visual was created using iTol^149^.

From inspection of fai’s resulting XLSX spreadsheet, zol’s parameters were adjusted to relax identity and coverage thresholds for assessing protein pairs for orthology prior to MCL clustering to 20% and 25%, respectively. Identical processing was performed for the full set of *epa* loci and *epa* loci from only species representative genomes. During the comprehensive processing of all high-quality *epa* loci identified, one instance was dropped during zol analysis despite meeting requirements because all CDS features in it were found near scaffold edges and, by default, such features are not used in zol to aid more accurate inference of ortholog groups and assessment of their sequence variation. A third run of zol was performed using identical settings and all the gene cluster instances but leveraging the dereplication and reinflation options to showcase how the combination of the options can reduce the runtime needed for comprehensive processing. For dereplication of gene clusters, alignment fraction was increased from the default of 95% to 99% and MCL was used for clustering to gather more resolute representative gene clusters. Major ortholog groups determined between the comprehensive and the dereplication + reinflation runs were found to be similarly conserved based on matching to known *epa* genes.

#### Phylogenetic assessment of glycosyltransferase orthology predictions

Proteins from ortholog groups determined by zol analysis of species representative genomes were extracted based on whether the ortholog group was annotated as featuring the keywords: “glycosyl” and “transferase” in Pfam protein domain annotations^150^. Two additional ortholog groups were included and featured the Pfam domain “Bacterial sugar transferase”, including *epaR*, which is also regarded as a glycosyltransferase^101^. The comprehensive set of glycosyltransferases were next aligned using MUSCLE with the default align mode^132^. Filtering of the alignment was next performed using trimal with options “-keepseqs -gt 0.9” to filter sites composed largely of gaps and further filtered for sequences which were composed of >10% gaps or ambiguous characters (“X”). IQ-TREE^151^ was used to construct a maximum-likelihood phylogeny with ModelFinder limited to the WAG and LG substitution models. The phylogeny was visualized using iTol^149^ with classifications for ortholog groups most closely matching *E. faecalis* V583 *epa* glycosyltransferases marked on leaves. Ortholog groups were assigned to specific *epa* gene designations based on sequence alignment of their consensus sequences to *E. faecalis* V583 *epa*-associated proteins. Best matching ortholog groups for each *E. faecalis* V583 *epa* glycosyltransferase were identified based on E-value.

### Large-scale evolutionary investigations of *epa* loci from *E. faecalis*

The full set of *epa* loci identified by fai in *E. faecalis* genomes were processed through zol requesting for retention of only complete instances that were also distant from scaffold edges. For projection of conservation, Tajima’s D, and sequence entropy statistics onto genes for the *epa* locus in *E. faecalis* V583, sequence alignment was used to identify the best matching ortholog groups based on E-value. For the identical transposases, EF2173 and EF2185, data from a common ortholog group was used for both.

#### Investigation of glycosyltransferase phylogenetic diversity

A similar phylogeny of glycosyltransferases was constructed for the *E. faecalis* analysis as was done for the investigation of *epa* glycosyltransferases across species representatives of *Enterococcus*. Glycosyltransferase ortholog groups were identified based on Pfam domains featuring the keywords “glycosyl transferase” or because they matched *epa* genes regarded as glycosyltransferases in prior studies^101^. To accommodate for the larger number of sequences: (i) only ortholog groups found in >1% of *epa* loci instances were regarded, (ii) MUSCLE^132^ super5 mode was used for alignment, and (iii) FastTree 2^127^ was used for approximate maximum-likelihood phylogeny construction. After trimal based filtering of sites, only sequences which featured greater than 20% gaps or ambiguous characters (“X”) were filtered to retain *epa*A in the final alignment prior to phylogeny construction.

## Abbreviations

BGC: Biosynthetic gene cluster
MGE: mobile-genetic element
Epa: Enterococcal polysaccharide antigen
CDS: coding sequence
ANI: average nucleotide identity
MAG: metagenome-assembled genome

## Declarations

### Ethics approval and consent to participate

Not applicable

### Consent for publication

Not applicable

### Availability of data and materials

All genomic and metagenomic datasets used for showcasing the application of fai and zol are publicly available on NCBI with accessions provided in Supplementary Table S11.

### Authors’ contributions

RS & LRK designed the bioinformatics toolkit. RS, KA, & LRK conceived of showcase applications. RS developed the software and performed all analyses. PQT performed assembly of metagenomes. RS & LRK performed the initial drafting of the manuscript. RS, PQT, CM, ALM, MSG, AME, KA, and LRK performed interpretation of results and revision of the manuscript. All authors read and approved the final manuscript.

## Acknowledgments

The authors are grateful to James Kosmopoulos, Dr. Caitlin Pepperell, Dr. Caitlin Sande, and Dr. Mary Hannah Swaney for feedback and assistance with data acquisition as well as Dr. Devon Ryan and Dr. Robert A. Petit III for assistance with incorporation of the suite into Bioconda.

