## Supplementary Text for "zol & fai: large-scale targeted detection and evolutionary investigation of gene clusters"

### **Differences between fai and cblaster**

fai is similar to cblaster^1^, another software for high-throughput identification of homologous gene clusters. However, each program has a different range of functionalities and options which gives them unique advantages depending on research objectives and use-cases (**Table S1**). Both cblaster and fai take as input a query gene cluster, provided as a set of proteins in FASTA or GenBank format, and use DIAMOND to rapidly search for homologous proteins in target genomes. In addition, for bacterial applications, fai can also take coordinates of a gene cluster along a reference genome. If query gene clusters are provided as a GenBank or coordinates along a reference genome, where gene order and direction information are preserved, fai can directly assess global syntenic similarity of candidate homologous gene clusters to the reference query gene cluster using a correlation based methodology^2^.

Unlike cblaster, fai performs a preliminary step to remove redundancy in query proteins using CD-HIT^3^ or, if multiple gene clusters are provided as a single query, the ortholog grouping algorithm used by zol. Bitscores are then used to assign the best matching non-redundant query protein clusters to coding sequences in target genomes. In contrast, cblaster searches for matches for each query protein independently and can thus assign multiple query proteins to the same coding sequence in the target genome. This can lead to unintended results when the query gene cluster involves homologous proteins, such as a BGC featuring recently duplicated non-ribosomal protein synthetases (NRPSs)^4^. Additionally, while cblaster and fai both rely on user-defined E-value cutoffs to identify homology, cblaster further allows filtering using percent identity and coverage cutoffs. A coverage cutoff option is not provided within fai because differences in gene-calling between the reference gene cluster and target genomes might result in a query protein not being fully represented by a predicted coding sequence (CDS) in target genomes (**Figure S1A**).

Another key difference between cblaster and fai is the approach taken to delineate candidate gene clusters. fai provides two different approaches for gene cluster delineation, one, referred to as the “Gene-Clumper” approach, which is similar to the approach taken by cblaster and simply groups together genes into the same cluster if they are separated by *M* or fewer CDS features, where *M* is by default set to 5. The other approach is based on a Hidden Markov Model (HMM) with tunable customizable transition and emission probabilities^5^ which can be used to first identify smaller sets of gene clusters and then aggregate those with the maximum CDS separator parameter described for the “Gene-Clumper” approach.

To aid selection of optimized parameters, cblaster provides an innovative analytical mode (gene neighborhood estimation; GNE) to infer the appropriate distance to allow between proteins homologous to the query to regard them as corresponding to the same gene cluster instance. Starting in v1.3.19, we introduced an option in zol to infer the optimal parameters for searching for additional instances of a gene cluster with fai, provided a handful of known homologous instances of the gene cluster as input. This mode can be requested via the argument “--select_fai_params_mode”.

Finally, cblaster, and the web-application CAGECAT^6^ - which runs cblaster, offer remote searching of BLAST databases, which fai does not support. In contrast, fai and prepTG, the software used for target genome database construction, feature unique options and capabilities ranging from integrated gene-calling or protein-mapping software to finding gene clusters split due to assembly fragmentation (**Figure S1B**).

### **Inference of fai searching parameters across homologous BGCs and phages**

To determine prior distributions for search parameters in fai for BGC and phage gene clusters, we applied zol to sets of homologous BGCs and phages with mode “--select_fai_params_mode” enabled (**Figure S3**). To gather sets of homologous phages, we downloaded viral genomes from NCBI using ncbi-acc-download (<https://github.com/kblin/ncbi-acc-download>) and partitioned them according to cluster categorizations in PhamClust^7^. To identify sets of homologous BGCs, we downloaded all characterized BGCs from the MIBiG database (v3.1)^8^ and clustered them into gene cluster families (GCFs) using BiG-SCAPE^9^ with “mix” mode requested to perform a comprehensive clustering. BGCs were partitioned based on their GCF classifications and, similar to homologous virus sets corresponding to distinct PhamClust clusters, were independently processed using zol.

### **Evolutionary-simulation based assessment of ortholog group inference by zol**

To directly assess the accuracy of zol to perform context-limited ortholog group inference relative to alternate software, we used evolutionary simulation of the *epa* locus from *E. faecalis* to derive a dataset where true orthologous relations can be inferred based on alignment to proteins from the original, “ancestral”, sequence.

The *epa* locus and its surrounding +/- 100 kb context from the *E. faecalis* V583 genome were extracted and a random tree to guide evolutionary simulation was constructed using the randomtree function from the ape library in R^10^. These were used as the primary inputs to run CoreSimul for evolutionary simulation^11^. CoreSimul was run using the Kimura two-parameter^12^ substitution model with the transition/transversion ratio set to 2^13^, the mutation rate of codon positions set to be equivalent (because our input included intergenic sequences), and the frequency of sequence gains and losses relative to the substitution rate set to 0.001 and 0.002^14^, respectively. After simulation, sequences were extracted and, together with the ancestral sequence, processed through prepTG to perform gene calling using pyrodigal^15^. Next, fai was used to search and extract only the *epa* locus (EF2164-EF2200) from the evolved sequences. The ‘--generate-plots’ flag was requested in fai to create schematics of how proteins from the unaltered, actual, *epa* locus of *E. faecalis* V583 aligned to corresponding fragmented or expanded genes in the evolved sequences (**Figure S7A**). Following visual inspection, coding-sequences in the evolved sequences were grouped into ortholog groups based on the protein from the unaltered locus (also the query gene cluster) that they best aligned to. Note, because a transposase in the locus is found in duplicate at 100% identity, two of the ortholog groups were merged into one.

The fai-extracted *epa* gene clusters in GenBank format were then processed through zol and clinker. For clinker, the report for pairs of matched proteins were clustered using single-linkage clustering with slclust (<https://sourceforge.net/projects/slclust/>) to identify sets of homologous proteins, to match the approach clinker takes internally^16^. Proteins from the GenBank files were also extracted and used as input for OrthoFinder. OrthoFinder and clinker were run using default parameters. Similarly, zol was also run using default parameters, but an additional run was also performed with modified parameters, whereby, based on reports generated by fai, we set the percent identity (default 30%) and coverage thresholds (default 50%) needed for pairs of proteins to be regarded as potential orthologs or in-paralogs to 0.

Comparisons of orthologs inferred by the different methods was performed directly against the inferred true orthologous relationships based on protein alignment (**Figure S7BCDE**). For each method, we assessed the number of true positives (ortholog pairs found in the truth set and inferred by the orthology inference method), false negatives (ortholog pairs found in the truth set but not by the orthology inference method), and false positives (ortholog pairs found by the orthology inference method but not in the truth set). Because it is possible for adjacent genes to be merged in evolved sequences, we also separately assessed the number of false positives discounting gene pairs which are direct neighbors in the locus. Ultimately, we found that zol-based orthology inference, with loosened identity and coverage thresholds, identified the greatest number of true positive ortholog pairs without incurring a substantial number of false positives. Coarse orthology inference by OrthoFinder (the first round of ortholog groups – equivalent to the output of OrthoFinder1^17^) and pairs of proteins belonging to the same single-linkage cluster of homologous proteins identified by clinker were both able to achieve the greatest recall of true positive orthologs; however, these methods also incurred thousands of false positive ortholog predictions. Refinement of ortholog groups in OrthoFinder (based on gene-to-species tree reconciliation described in OrthoFinder2^18^) performed similar to zol with loosened criteria; however, zol found more true positives and fewer false negatives. In addition, zol with default identity and coverage thresholds for orthology inference reported 0 false negative ortholog pairs.

### **Dereplication can impact taxa-wide inferences of selection-informative statistics**

The impacts of zol’s dereplication related approaches on the calculation of evolutionary statistics were assessed using 1,232 instances of *epa* from *E. faecalis* that were high-quality (<10% of bases ambiguous) and not near scaffold edges. Specifically, we compared values for evolutionary statistics calculated by running zol with: (i) no dereplication, (ii) only instances of the *epa* locus from genome-wide representatives selected by dRep with a cutoff of 99% average nucleotide identity (ANI)^19,20^, (iii) gene cluster dereplication with cutoffs of 99% ANI and 99% aligned fraction (AF), (iv) gene cluster dereplication with cutoffs of 97% ANI and 95% AF, and (v) gene cluster dereplication with cutoffs of 99% ANI and 99% aligned fraction followed by ortholog group reinflation. Reinflation, which can optionally be requested together with dereplication, allows for expansion of ortholog groups determined from representative gene clusters to include proteins from gene clusters which were deemed redundant during dereplication. The purpose of reinflation is to alleviate time and computational resources required for comprehensive investigations of a set of gene clusters. For each zol analysis, the best matching ortholog group for *epa* proteins EF2164-EF2200 (not including EF2173 and EF2185-EF2188 because they represent transposases or interrupted genes in *E. faecalis* V583^21^) were identified based on E-value. Evolutionary statistics were then compared between corresponding ortholog groups mapping to the same *epa* protein. Ortholog groups which were designated as non-single copy by zol were not accounted for in comparisons. To assess the reproducibility of values for evolutionary statistics between the five different zol runs the concordance correlation coefficient was computed^22^.

Estimates of evolutionary and genomic statistics for analogous ortholog groups were found to be robust for some metrics, but not for all (**Figure S9, S10**). For instance, dereplication can impact selection informative statistics, such as FUBAR-based^23^ inference of the proportion of total sites under selection that are specifically under positive selection. While such phylogeny informed statistics should be robust to the effects of dereplication, it is possible that dereplication alters the gene phylogeny for an ortholog group resulting in slight differences in posterior distributions of the non-synonymous and synonymous substitution rates inferred by FUBAR. The values for such statistics still trend for analogous ortholog groups across different runs of zol where dereplication is used in different ways or not performed (**Figure S10**). Other statistics were relatively robust to the application of dereplication, such as Tajima’s D. In addition, provided the discordance between statistics following dereplication using different thresholds for clustering of gene cluster instances, such thresholds should be carefully considered during interpretation of evolutionary trends.

Of practical importance, skipping dereplication produced highly similar statistical values to a zol analysis where dereplication was performed followed by reinflation, simulating comprehensive processing of gene clusters. The latter approach is much faster due to avoiding extensive all-vs-all alignment needed to determine ortholog groups across all gene cluster instances; instead limiting ortholog group delineation to be performed across only distinct representative gene cluster instances (**Figure S8**).

### **Parameters used in fai and zol for showcase applications**

Parameters for fai and zol are reported here for major analyses highlighted in the manuscript, parameters and command files for additional fai and zol analyses mentioned in the manuscript can be found at: <https://github.com/Kalan-Lab/Salamzade_etal_zol>.

**Parameters used for** **“Application of fai and zol to identify phages within metagenomes”**

**Parameters used for fai analysis:**

Query gene-cluster GenBanks: ['Query_Phage/Ga0485157_0000450.gbk', 'Query_Phage/Ga0485158_0000434.gbk']

Reference genome for gene cluster: None

Reference scaffold for gene cluster: None

Reference start coordinate of gene cluster: None

Reference end coordinate of gene cluster: None

Protein queries multi-FASTA file: None

Single protein query FASTA file: None

Target genomes prepared directory by prepTG: prepTG_DB/

Output directory: /home/salamzade/zol_development/showcase_examples_redo/Phage_in_Lake_MGs/fai_Results/

Species Tree: None

Run in draft-assembly mode?: False

Filter for paralogous/gene-content overlapping segments: True

Use prodigal-gv instead of pyrodigal to perform gene calling for query region in reference genome: False

General E-value cutoff for detection of protein homologs in genome: 1e-10

Minimum proportion of distinct query proteins needed: 0.5

FASTA file with key proteins to consider: None

E-values for key proteins to be considered as high-supporting evidence for gene cluster presence: 1e-20

Minimum proportion of distinct key protein queries needed: 0.0

Syntenic correlation to known instance threshold needed: 0.0

Maximum distance in between candidate gene-cluster segments to perform merging: 5

Base pair for flanking context to extract: 1000

Emission probability of gene being in gene-cluster state with homologous hit to gene-cluster: 0.95

Emission probability of gene being in background state with homologous hit to gene-cluster: 0.2

Probability for gene-cluster to gene-cluster transition in HMM: 0.9

Probability for background to background transition in HMM: 0.9

Perform plotting?: False

DIAMOND sensitivity: very-sensitive

Delineation mode: GENE-CLUMPER

Phylo-heatmap PDF length: 7

Phylo-heatmap PDF width: 14

Number of CPUs requested: 20

Maximum memory in GB: None

Clean Up Heavy Files?: False

**Parameters used for zol analysis:**

Input directory with loci GenBanks: /home/salamzade/zol_development/showcase_examples_redo/Phage_in_Lake_MGs/fai_Results/Final_Results/Homologous_Gene_Cluster_GenBanks/

Output directory: /home/salamzade/zol_development/showcase_examples_redo/Phage_in_Lake_MGs/zol_Results/

Select fai parameters mode?: False

Perform iterative CD-HIT based orthogrouping instead of default InParanoid-based approach?: False

CD-HIT parameters for orthogrouping (assuming approach requested): -c 0.5 -aL 0.25 -aS 0.5 -n 3 -M 4000

Ortholog group finding identity threshold: 30.0

Ortholog group finding coverage threshold: 50.0

Ortholog group finding E-value threshold: 0.001

Use super5 mode in MUSCLE alignments?: False

Run FUBAR selection analyses?: False

Skip GARD partitioning by recombination breakpoints?: False

Focal GenBanks listing: None

Comparator GenBanks listing: None

Filter low quality gene clusters?: False

Filter draft/incomplete gene clusters?: False

Only compute orthologs and stop?: False

Perform broad level estimation of ortholog group conservation if dereplication requested?: False

Comprehensive reporting of evolutionary statistics, including for non-single copy ortholog groups: False

Rename locus tags?: False

Use CDS features with attribute near_scaffold_edge=True.: True

Perform Dereplication?: False

Perform reinflation?: False

Reinflation CD-HIT parameters: -c 0.98 -aL 0.95 -aS 0.95 -n 5 -M 4000

Dereplication identity threshold: 99.0

Dereplication coverage threshold: 95.0

Dereplication clustering method / MCL inflation: None

Custom annotation database: None

Refine gene calling using the custom annotation database: False

Plot height: 7

Plot width: 14

Use full GenBank labels?: False

Number of CPUs requested: 20

Maximum memory in GB: None

**Parameters used for “Microevolutionary investigations of leporin and aflatoxin BGCs in *Aspergillus flavus*”**

**Parameters used for fai analysis of leporin BGC:**

Query gene-cluster GenBanks: ['BGC0001445.gbk']

Reference genome for gene cluster: None

Reference scaffold for gene cluster: None

Reference start coordinate of gene cluster: None

Reference end coordinate of gene cluster: None

Protein queries multi-FASTA file: None

Single protein query FASTA file: None

Target genomes prepared directory by prepTG: ../prepTG_DB/

Output directory: /home/salamzade/zol_development/showcase_examples_redo/Aflavus_aflatoxin_and_leporinB/leporin/fai_Results/

Species Tree: None

Run in draft-assembly mode?: True

Filter for paralogous/gene-content overlapping segments: True

Use prodigal-gv instead of pyrodigal to perform gene calling for query region in reference genome: False

General E-value cutoff for detection of protein homologs in genome: 1e-10

Minimum proportion of distinct query proteins needed: 0.5

FASTA file with key proteins to consider: key_proteins.faa

E-values for key proteins to be considered as high-supporting evidence for gene cluster presence: 1e-20

Minimum proportion of distinct key protein queries needed: 1.0

Syntenic correlation to known instance threshold needed: 0.6

Maximum distance in between candidate gene-cluster segments to perform merging: 3

Base pair for flanking context to extract: 1000

Emission probability of gene being in gene-cluster state with homologous hit to gene-cluster: 0.95

Emission probability of gene being in background state with homologous hit to gene-cluster: 0.2

Probability for gene-cluster to gene-cluster transition in HMM: 0.9

Probability for background to background transition in HMM: 0.9

Perform plotting?: True

DIAMOND sensitivity: very-sensitive

Delineation mode: GENE-CLUMPER

Phylo-heatmap PDF length: 7

Phylo-heatmap PDF width: 14

Number of CPUs requested: 20

Maximum memory in GB: None

Clean Up Heavy Files?: False

**Parameters used for zol analysis of leporin BGC:**

Input directory with loci GenBanks: /home/salamzade/zol_development/showcase_examples_redo/Aflavus_aflatoxin_and_leporinB/leporin/fai_Results/Final_Results/Homologous_Gene_Cluster_GenBanks/

Output directory: /home/salamzade/zol_development/showcase_examples_redo/Aflavus_aflatoxin_and_leporinB/leporin/zol_Results/

Select fai parameters mode?: False

Perform iterative CD-HIT based orthogrouping instead of default InParanoid-based approach?: False

CD-HIT parameters for orthogrouping (assuming approach requested): -c 0.5 -aL 0.25 -aS 0.5 -n 3 -M 4000

Ortholog group finding identity threshold: 30.0

Ortholog group finding coverage threshold: 50.0

Ortholog group finding E-value threshold: 0.001

Use super5 mode in MUSCLE alignments?: False

Run FUBAR selection analyses?: False

Skip GARD partitioning by recombination breakpoints?: False

Focal GenBanks listing: PopB_GCs.txt

Comparator GenBanks listing: None

Filter low quality gene clusters?: True

Filter draft/incomplete gene clusters?: True

Only compute orthologs and stop?: False

Perform broad level estimation of ortholog group conservation if dereplication requested?: False

Comprehensive reporting of evolutionary statistics, including for non-single copy ortholog groups: False

Rename locus tags?: False

Use CDS features with attribute near_scaffold_edge=True.: False

Perform Dereplication?: False

Perform reinflation?: False

Reinflation CD-HIT parameters: -c 0.98 -aL 0.95 -aS 0.95 -n 5 -M 4000

Dereplication identity threshold: 99.0

Dereplication coverage threshold: 95.0

Dereplication clustering method / MCL inflation: None

Custom annotation database: BGC0001445.faa

Refine gene calling using the custom annotation database: False

Plot height: 7

Plot width: 14

Use full GenBank labels?: False

Number of CPUs requested: 20

Maximum memory in GB: None

**Parameters used for fai analysis of aflatoxin BGC:**

Query gene-cluster GenBanks: ['BGC0000008.gbk']

Reference genome for gene cluster: None

Reference scaffold for gene cluster: None

Reference start coordinate of gene cluster: None

Reference end coordinate of gene cluster: None

Protein queries multi-FASTA file: None

Single protein query FASTA file: None

Target genomes prepared directory by prepTG: ../prepTG_DB/

Output directory: /home/salamzade/zol_development/showcase_examples_redo/Aflavus_aflatoxin_and_leporinB/aflatoxin/fai_Results/

Species Tree: None

Run in draft-assembly mode?: True

Filter for paralogous/gene-content overlapping segments: True

Use prodigal-gv instead of pyrodigal to perform gene calling for query region in reference genome: False

General E-value cutoff for detection of protein homologs in genome: 1e-10

Minimum proportion of distinct query proteins needed: 0.5

FASTA file with key proteins to consider: key_proteins.faa

E-values for key proteins to be considered as high-supporting evidence for gene cluster presence: 1e-20

Minimum proportion of distinct key protein queries needed: 1.0

Syntenic correlation to known instance threshold needed: 0.6

Maximum distance in between candidate gene-cluster segments to perform merging: 3

Base pair for flanking context to extract: 1000

Emission probability of gene being in gene-cluster state with homologous hit to gene-cluster: 0.95

Emission probability of gene being in background state with homologous hit to gene-cluster: 0.2

Probability for gene-cluster to gene-cluster transition in HMM: 0.9

Probability for background to background transition in HMM: 0.9

Perform plotting?: True

DIAMOND sensitivity: very-sensitive

Delineation mode: GENE-CLUMPER

Phylo-heatmap PDF length: 7

Phylo-heatmap PDF width: 14

Number of CPUs requested: 20

Maximum memory in GB: None

Clean Up Heavy Files?: False

**Parameters used for zol analysis of aflatoxin BGC:**

Input directory with loci GenBanks: /home/salamzade/zol_development/showcase_examples_redo/Aflavus_aflatoxin_and_leporinB/aflatoxin/fai_Results/Final_Results/Homologous_Gene_Cluster_GenBanks/

Output directory: /home/salamzade/zol_development/showcase_examples_redo/Aflavus_aflatoxin_and_leporinB/aflatoxin/zol_Results/

Select fai parameters mode?: False

Perform iterative CD-HIT based orthogrouping instead of default InParanoid-based approach?: False

CD-HIT parameters for orthogrouping (assuming approach requested): -c 0.5 -aL 0.25 -aS 0.5 -n 3 -M 4000

Ortholog group finding identity threshold: 30.0

Ortholog group finding coverage threshold: 50.0

Ortholog group finding E-value threshold: 0.001

Use super5 mode in MUSCLE alignments?: False

Run FUBAR selection analyses?: False

Skip GARD partitioning by recombination breakpoints?: False

Focal GenBanks listing: None

Comparator GenBanks listing: None

Filter low quality gene clusters?: True

Filter draft/incomplete gene clusters?: True

Only compute orthologs and stop?: False

Perform broad level estimation of ortholog group conservation if dereplication requested?: False

Comprehensive reporting of evolutionary statistics, including for non-single copy ortholog groups: False

Rename locus tags?: False

Use CDS features with attribute near_scaffold_edge=True.: False

Perform Dereplication?: False

Perform reinflation?: False

Reinflation CD-HIT parameters: -c 0.98 -aL 0.95 -aS 0.95 -n 5 -M 4000

Dereplication identity threshold: 99.0

Dereplication coverage threshold: 95.0

Dereplication clustering method / MCL inflation: None

Custom annotation database: BGC0000008.faa

Refine gene calling using the custom annotation database: False

Plot height: 7

Plot width: 14

Use full GenBank labels?: False

Number of CPUs requested: 20

Maximum memory in GB: None

**Parameters used for “Evolutionary investigations of the *epa* locus across *Enterococcus*”**

**Parameters used for fai analysis:**

Query gene-cluster GenBanks: None

Reference genome for gene cluster: GCF_000007785.1_ASM778v1_genomic.fna

Reference scaffold for gene cluster: NC_004668.1

Reference start coordinate of gene cluster: 2071671

Reference end coordinate of gene cluster: 2115174

Protein queries multi-FASTA file: None

Single protein query FASTA file: None

Target genomes prepared directory by prepTG: prepTG_DB/

Output directory: /home/salamzade/zol_development/showcase_examples_redo/Enterococcus_Epa/fai_Results/

Species Tree: None

Run in draft-assembly mode?: True

Filter for paralogous/gene-content overlapping segments: True

Use prodigal-gv instead of pyrodigal to perform gene calling for query region in reference genome: False

General E-value cutoff for detection of protein homologs in genome: 1e-10

Minimum proportion of distinct query proteins needed: 0.1

FASTA file with key proteins to consider: key_proteins.faa

E-values for key proteins to be considered as high-supporting evidence for gene cluster presence: 1e-10

Minimum proportion of distinct key protein queries needed: 0.5

Syntenic correlation to known instance threshold needed: 0.0

Maximum distance in between candidate gene-cluster segments to perform merging: 5

Base pair for flanking context to extract: 20000

Emission probability of gene being in gene-cluster state with homologous hit to gene-cluster: 0.95

Emission probability of gene being in background state with homologous hit to gene-cluster: 0.2

Probability for gene-cluster to gene-cluster transition in HMM: 0.9

Probability for background to background transition in HMM: 0.9

Perform plotting?: False

DIAMOND sensitivity: very-sensitive

Delineation mode: GENE-CLUMPER

Phylo-heatmap PDF length: 7

Phylo-heatmap PDF width: 14

Number of CPUs requested: 30

Maximum memory in GB: None

Clean Up Heavy Files?: False

**Parameters used for *Enterococcus* comprehensive zol analysis:**

Input directory with loci GenBanks: /home/salamzade/zol_development/showcase_examples_redo/Enterococcus_Epa/fai_Results/Final_Results/Homologous_Gene_Cluster_GenBanks/

Output directory: /home/salamzade/zol_development/showcase_examples_redo/Enterococcus_Epa/OneSpeciesOneRep_Phylo_Heatmap_and_GT_Phylogeny/zol_Results_Full/

Select fai parameters mode?: False

Perform iterative CD-HIT based orthogrouping instead of default InParanoid-based approach?: False

CD-HIT parameters for orthogrouping (assuming approach requested): -c 0.5 -aL 0.25 -aS 0.5 -n 3 -M 4000

Ortholog group finding identity threshold: 20.0

Ortholog group finding coverage threshold: 25.0

Ortholog group finding E-value threshold: 0.001

Use super5 mode in MUSCLE alignments?: True

Run FUBAR selection analyses?: False

Skip GARD partitioning by recombination breakpoints?: False

Focal GenBanks listing: None

Comparator GenBanks listing: None

Filter low quality gene clusters?: True

Filter draft/incomplete gene clusters?: False

Only compute orthologs and stop?: False

Perform broad level estimation of ortholog group conservation if dereplication requested?: False

Comprehensive reporting of evolutionary statistics, including for non-single copy ortholog groups: False

Rename locus tags?: False

Use CDS features with attribute near_scaffold_edge=True.: False

Perform Dereplication?: False

Perform reinflation?: False

Reinflation CD-HIT parameters: -c 0.98 -aL 0.95 -aS 0.95 -n 5 -M 4000

Dereplication identity threshold: 99.0

Dereplication coverage threshold: 95.0

Dereplication clustering method / MCL inflation: None

Custom annotation database: ../epa_prots_for_custom_annotation.faa

Refine gene calling using the custom annotation database: False

Plot height: 7

Plot width: 14

Use full GenBank labels?: False

Number of CPUs requested: 30

Maximum memory in GB: None

**Parameters used for *Enterococcus* species representative zol analysis:**

Input directory with loci GenBanks: /home/salamzade/zol_development/showcase_examples_redo/Enterococcus_Epa/OneSpeciesOneRep_Phylo_Heatmap_and_GT_Phylogeny/Rep_GCs/

Output directory: /home/salamzade/zol_development/showcase_examples_redo/Enterococcus_Epa/OneSpeciesOneRep_Phylo_Heatmap_and_GT_Phylogeny/zol_Results/

Select fai parameters mode?: False

Perform iterative CD-HIT based orthogrouping instead of default InParanoid-based approach?: False

CD-HIT parameters for orthogrouping (assuming approach requested): -c 0.5 -aL 0.25 -aS 0.5 -n 3 -M 4000

Ortholog group finding identity threshold: 20.0

Ortholog group finding coverage threshold: 25.0

Ortholog group finding E-value threshold: 0.001

Use super5 mode in MUSCLE alignments?: True

Run FUBAR selection analyses?: False

Skip GARD partitioning by recombination breakpoints?: False

Focal GenBanks listing: None

Comparator GenBanks listing: None

Filter low quality gene clusters?: True

Filter draft/incomplete gene clusters?: False

Only compute orthologs and stop?: False

Perform broad level estimation of ortholog group conservation if dereplication requested?: False

Comprehensive reporting of evolutionary statistics, including for non-single copy ortholog groups: False

Rename locus tags?: False

Use CDS features with attribute near_scaffold_edge=True.: False

Perform Dereplication?: False

Perform reinflation?: False

Reinflation CD-HIT parameters: -c 0.98 -aL 0.95 -aS 0.95 -n 5 -M 4000

Dereplication identity threshold: 99.0

Dereplication coverage threshold: 95.0

Dereplication clustering method / MCL inflation: None

Custom annotation database: ../epa_prots_for_custom_annotation.faa

Refine gene calling using the custom annotation database: False

Plot height: 7

Plot width: 14

Use full GenBank labels?: False

Number of CPUs requested: 30

Maximum memory in GB: None

**Parameters used for comprehensive *E. faecalis* zol analysis:**

Input directory with loci GenBanks: /home/salamzade/zol_development/showcase_examples_redo/Enterococcus_Epa/Efaecalis_GCs/

Output directory: /home/salamzade/zol_development/showcase_examples_redo/Enterococcus_Epa/Benchmarking_Dereplication/zol_Results/

Select fai parameters mode?: False

Perform iterative CD-HIT based orthogrouping instead of default InParanoid-based approach?: False

CD-HIT parameters for orthogrouping (assuming approach requested): -c 0.5 -aL 0.25 -aS 0.5 -n 3 -M 4000

Ortholog group finding identity threshold: 30.0

Ortholog group finding coverage threshold: 50.0

Ortholog group finding E-value threshold: 0.001

Use super5 mode in MUSCLE alignments?: True

Run FUBAR selection analyses?: True

Skip GARD partitioning by recombination breakpoints?: True

Focal GenBanks listing: None

Comparator GenBanks listing: None

Filter low quality gene clusters?: True

Filter draft/incomplete gene clusters?: True

Only compute orthologs and stop?: False

Perform broad level estimation of ortholog group conservation if dereplication requested?: False

Comprehensive reporting of evolutionary statistics, including for non-single copy ortholog groups: False

Rename locus tags?: False

Use CDS features with attribute near_scaffold_edge=True.: False

Perform Dereplication?: False

Perform reinflation?: False

Reinflation CD-HIT parameters: -c 0.98 -aL 0.95 -aS 0.95 -n 5 -M 4000

Dereplication identity threshold: 99.0

Dereplication coverage threshold: 95.0

Dereplication clustering method / MCL inflation: None

Custom annotation database: epa_prots_for_custom_annotation.faa

Refine gene calling using the custom annotation database: False

Plot height: 7

Plot width: 14

Use full GenBank labels?: False

Number of CPUs requested: 20

Maximum memory in GB: None

**Parameters used for comprehensive *Enterococcus* zol analysis with dereplication + reinflation:**

Input directory with loci GenBanks: /home/salamzade/zol_development/showcase_examples_redo/Enterococcus_Epa/fai_Results/Final_Results/Homologous_Gene_Cluster_GenBanks/

Output directory: /home/salamzade/zol_development/showcase_examples_redo/Enterococcus_Epa/OneSpeciesOneRep_Phylo_Heatmap_and_GT_Phylogeny/zol_Results_Full_DerepReinflate_DT99_DC99_DI2/

Select fai parameters mode?: False

Perform iterative CD-HIT based orthogrouping instead of default InParanoid-based approach?: False

CD-HIT parameters for orthogrouping (assuming approach requested): -c 0.5 -aL 0.25 -aS 0.5 -n 3 -M 4000

Ortholog group finding identity threshold: 20.0

Ortholog group finding coverage threshold: 25.0

Ortholog group finding E-value threshold: 0.001

Use super5 mode in MUSCLE alignments?: True

Run FUBAR selection analyses?: False

Skip GARD partitioning by recombination breakpoints?: False

Focal GenBanks listing: None

Comparator GenBanks listing: None

Filter low quality gene clusters?: True

Filter draft/incomplete gene clusters?: False

Only compute orthologs and stop?: False

Perform broad level estimation of ortholog group conservation if dereplication requested?: False

Comprehensive reporting of evolutionary statistics, including for non-single copy ortholog groups: False

Rename locus tags?: False

Use CDS features with attribute near_scaffold_edge=True.: False

Perform Dereplication?: True

Perform reinflation?: True

Reinflation CD-HIT parameters: -c 0.98 -aL 0.95 -aS 0.95 -n 5 -M 4000

Dereplication identity threshold: 99.0

Dereplication coverage threshold: 99.0

Dereplication clustering method / MCL inflation: 2.0

Custom annotation database: ../epa_prots_for_custom_annotation.faa

Refine gene calling using the custom annotation database: False

Plot height: 7

Plot width: 14

Use full GenBank labels?: False

Number of CPUs requested: 30

Maximum memory in GB: None

### **Supplementary text references**

1.	Gilchrist, C. L. M. *et al.* Cblaster: A remote search tool for rapid identification and visualization of homologous gene clusters. *Bioinformatics Advances* **1**, (2021).

2.	Salamzade, R. *et al.* Evolutionary investigations of the biosynthetic diversity in the skin microbiome using lsaBGC. *Microb Genom* **9**, (2023).

3.	Huang, Y., Niu, B., Gao, Y., Fu, L. & Li, W. CD-HIT Suite: a web server for clustering and comparing biological sequences. *Bioinformatics* **26**, 680–682 (2010).

4.	Medema, M. H., Cimermancic, P., Sali, A., Takano, E. & Fischbach, M. A. A systematic computational analysis of biosynthetic gene cluster evolution: lessons for engineering biosynthesis. *PLoS Comput. Biol.* **10**, e1004016 (2014).

5.	Schreiber, J. Pomegranate: fast and flexible probabilistic modeling in python. *J. Mach. Learn. Res.* (2017).
