## Supplementary Figures S1-S10 for "zol & fai: large-scale targeted detection and evolutionary investigation of gene clusters"

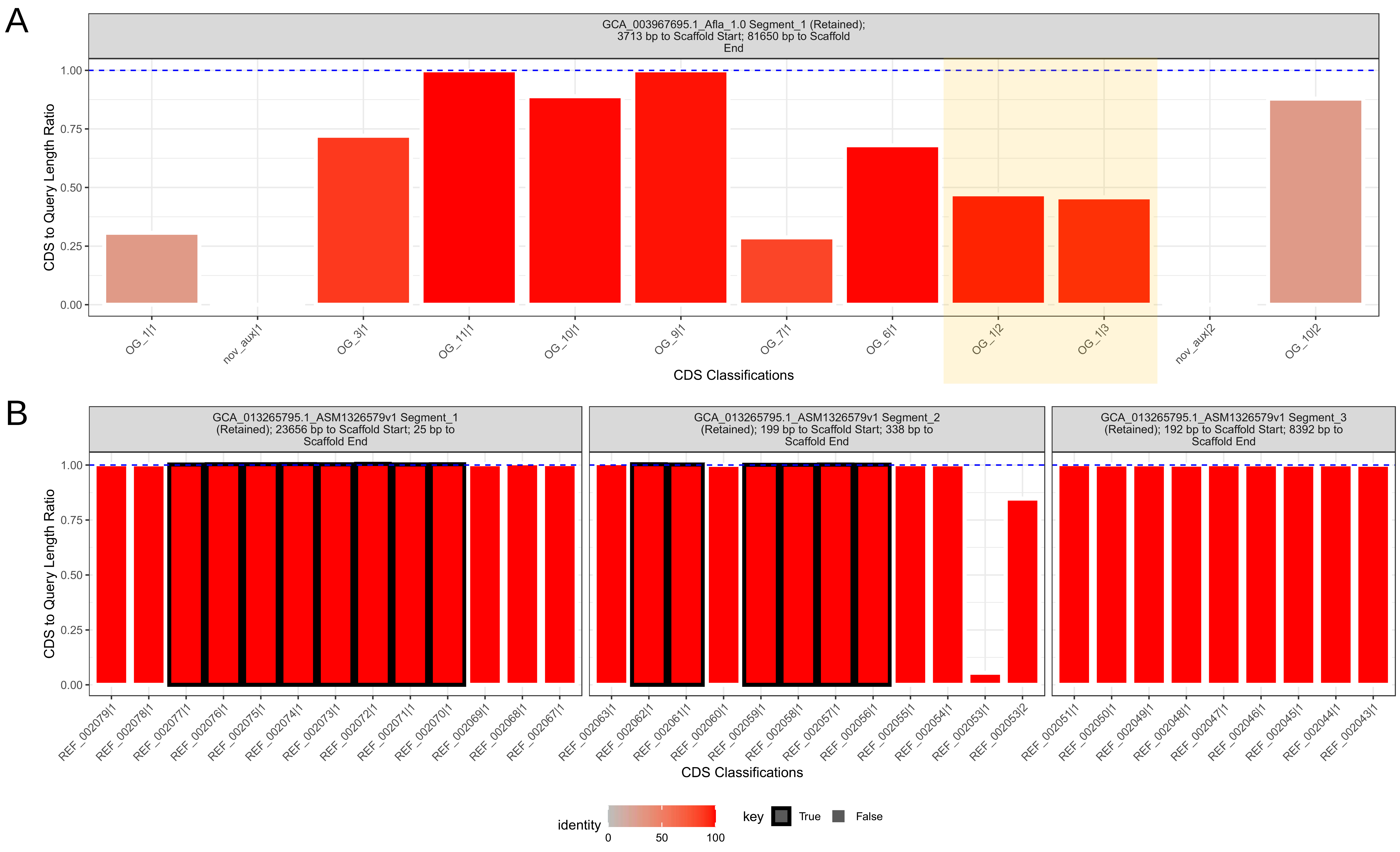


**Figure S1: Example illustrations for assessing quality of homologous gene-clusters produced by fai. A**) Gene calling or frame-shift differences between the query gene-cluster and coding-sequence predictions in the target genome have resulted in a discrepancy for OG_1 (highlighted) from the query being regarded as two separate coding-sequences in the target genome. **B**) Three candidate gene-cluster segments located near scaffold edges which match the query gene-cluster and meet the thresholds needed for detection as requested in fai in aggregate.

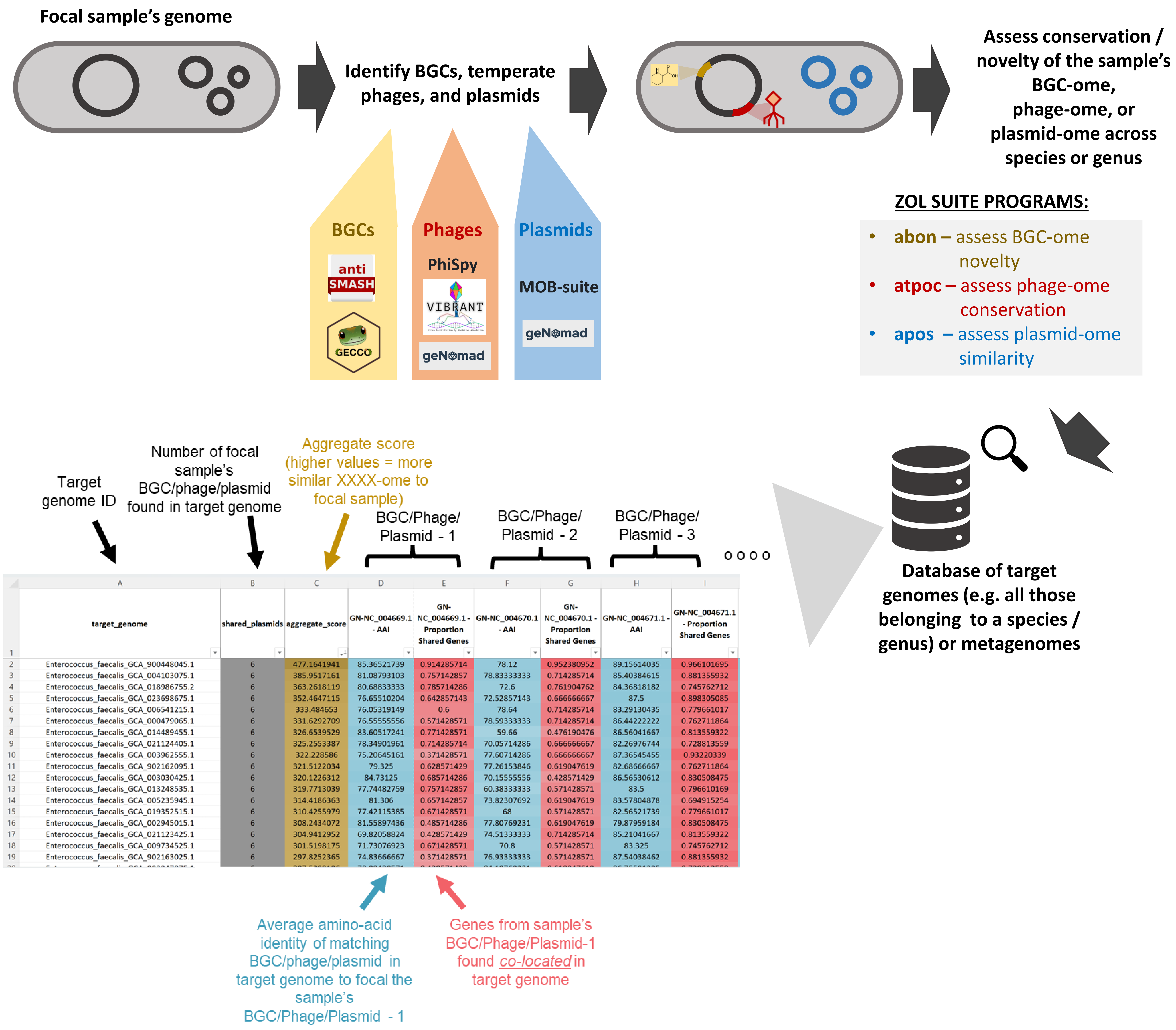


**Figure S2: Overview of abon, atpoc, and apos.** The zol suite includes tools abon, atpoc, and apos which are wrappers of fai and simplify checking whether a focal strain features novel BGCs, phages, and plasmids or assessing their conservation across a target database of genomes.

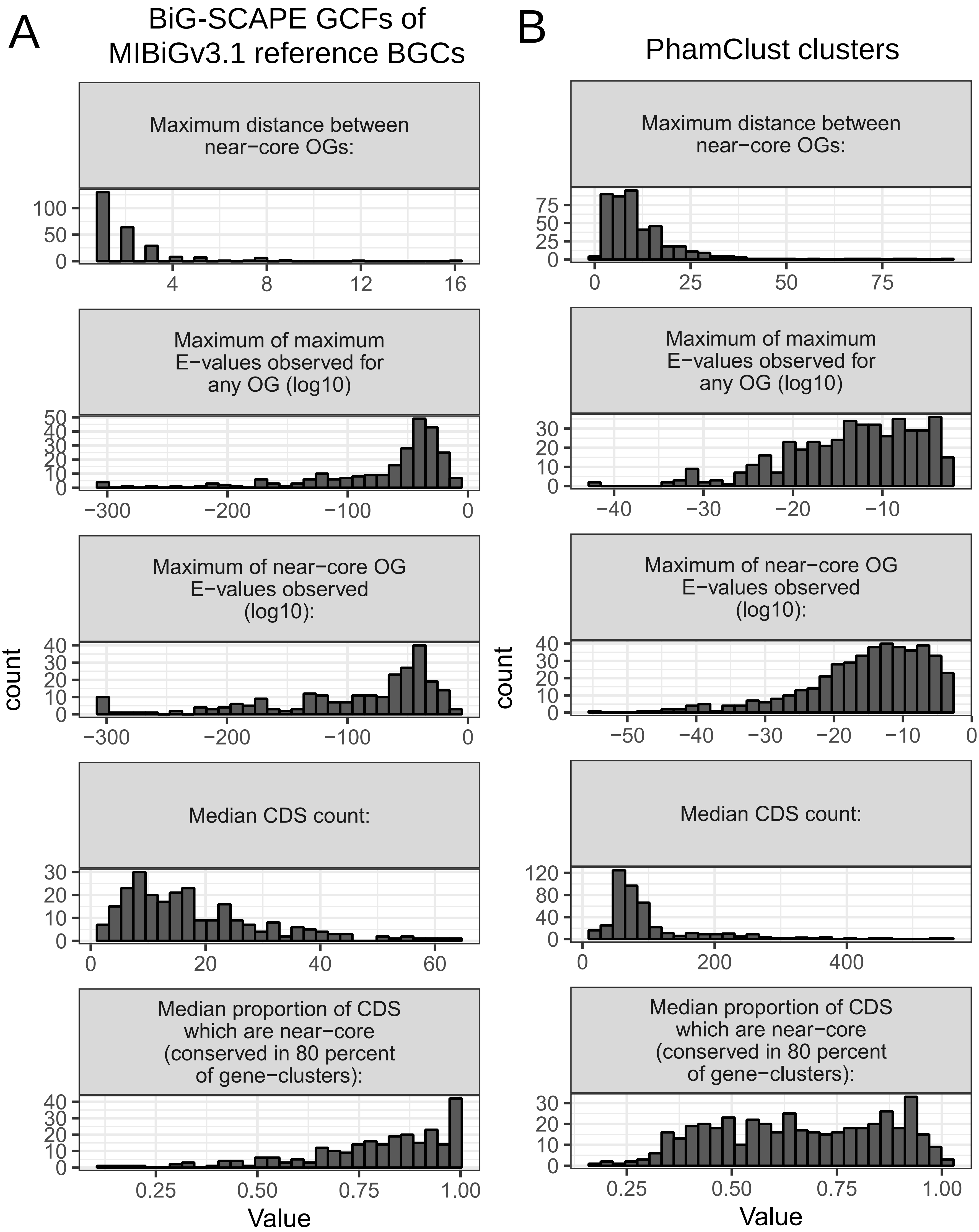


**Figure S3: Investigation of similarity statistics for BGC and phage clusters.** Distributions of intra-cluster similarity metrics which are informative for selection of parameters in fai are provided for **A**) GCFs of characterized BGCs from MIBiG v3.1 and **B**) PhamClust phage clusters.

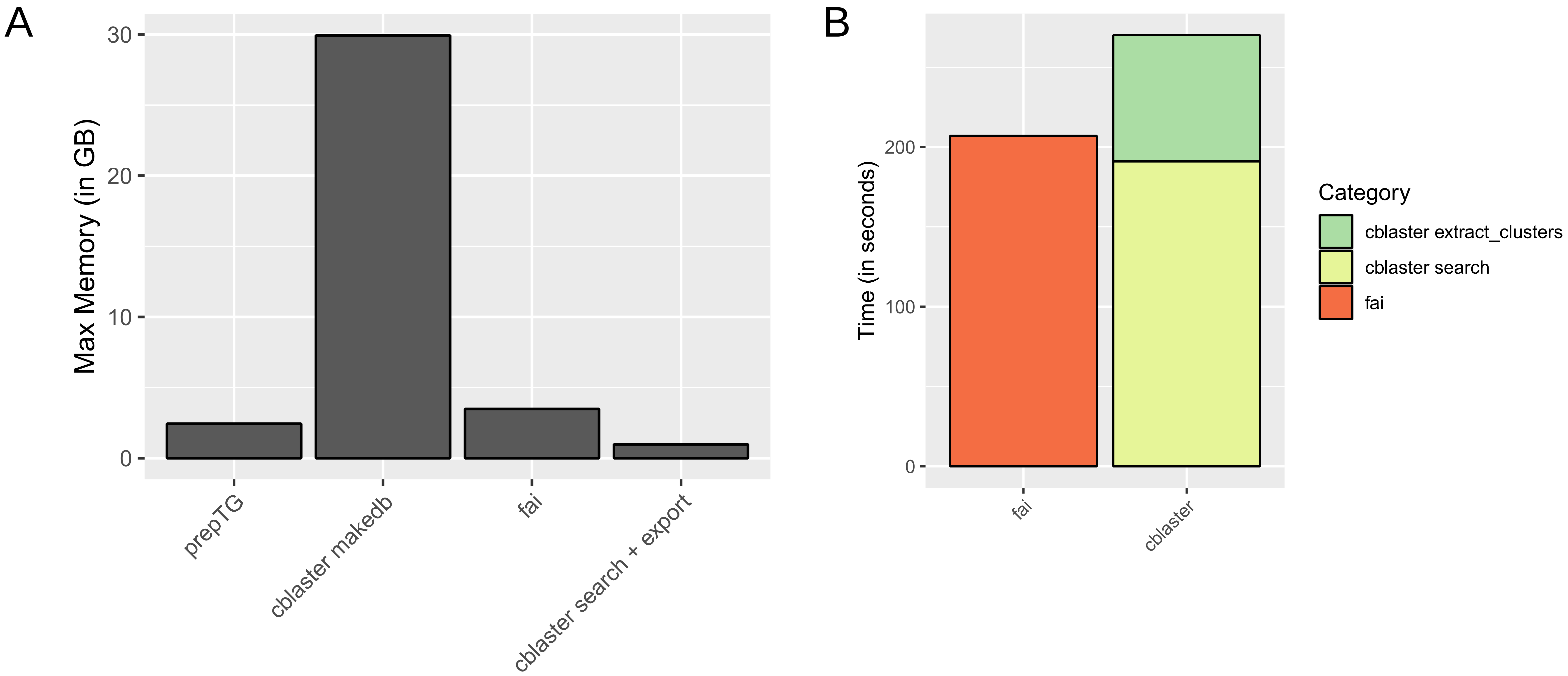


**Figure S4: Memory consumption and runtime comparisons between fai and cblaster for metagenomic applications. A)** The memory consumption of different commands from the zol and cblaster suites for preparing and searching metagenomic assemblies for gene clusters. **B**) The runtime for searching and extracting gene clusters in GenBank format using fai and cblaster.

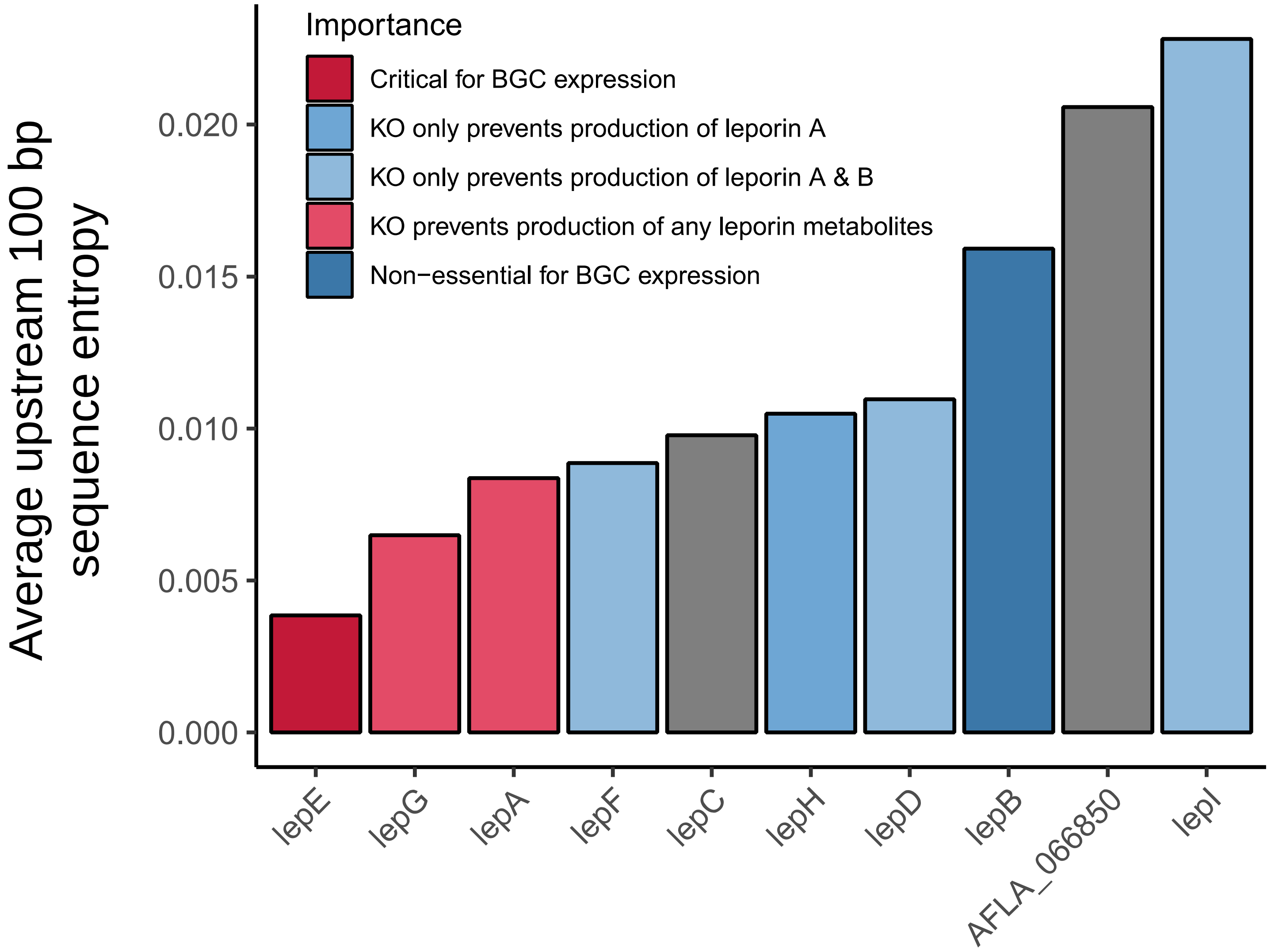


**Figure S5: Conservation in the upstream regions of coding sequences of genes in the leporin BGC.** The average entropy of the 100 bp upstream regions is shown for each of the genes from the leporin BGC. Coloring of the bars corresponds to effects on BGC expression (for *lepE* and *lepB*) or metabolite production (using a mutant with overexpression of *lepE*) when genes were knocked out as determined by Cary *et al.* 2015.

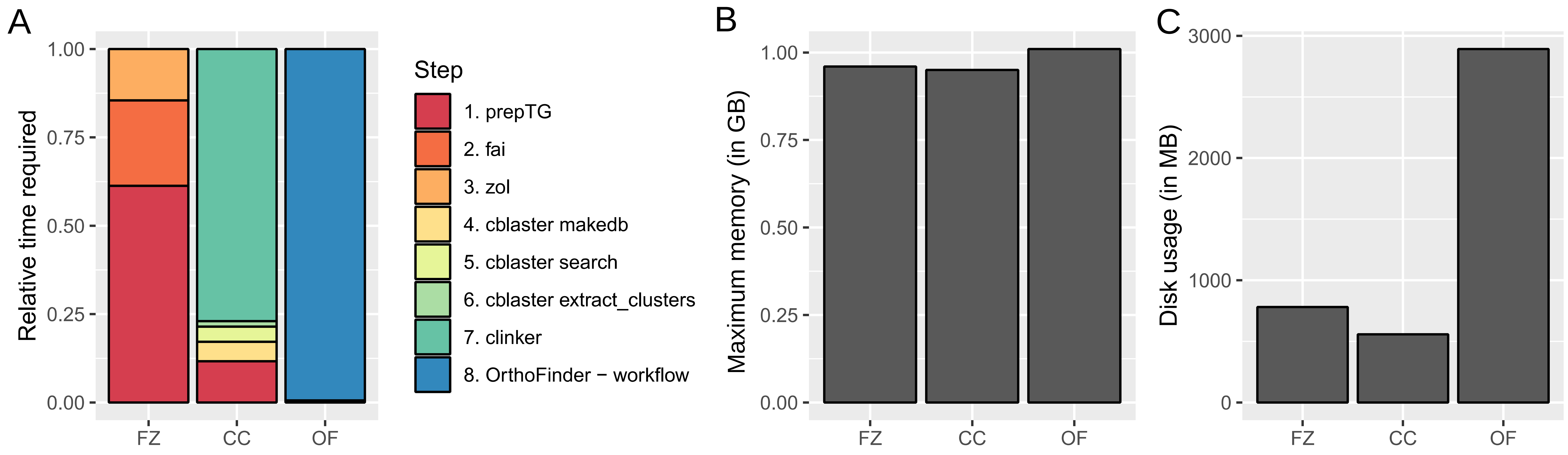


**Figure S6: Comparisons between fai & zol to alternate methods for inferring orthologs/homologs. A)** Breakdown of relative time to completion for the three different methods for interring orthologs/homologs by individual steps. An overview of the **B**) maximum memory usage in gigabytes and **C**) the final disk space taken by the three different approaches.

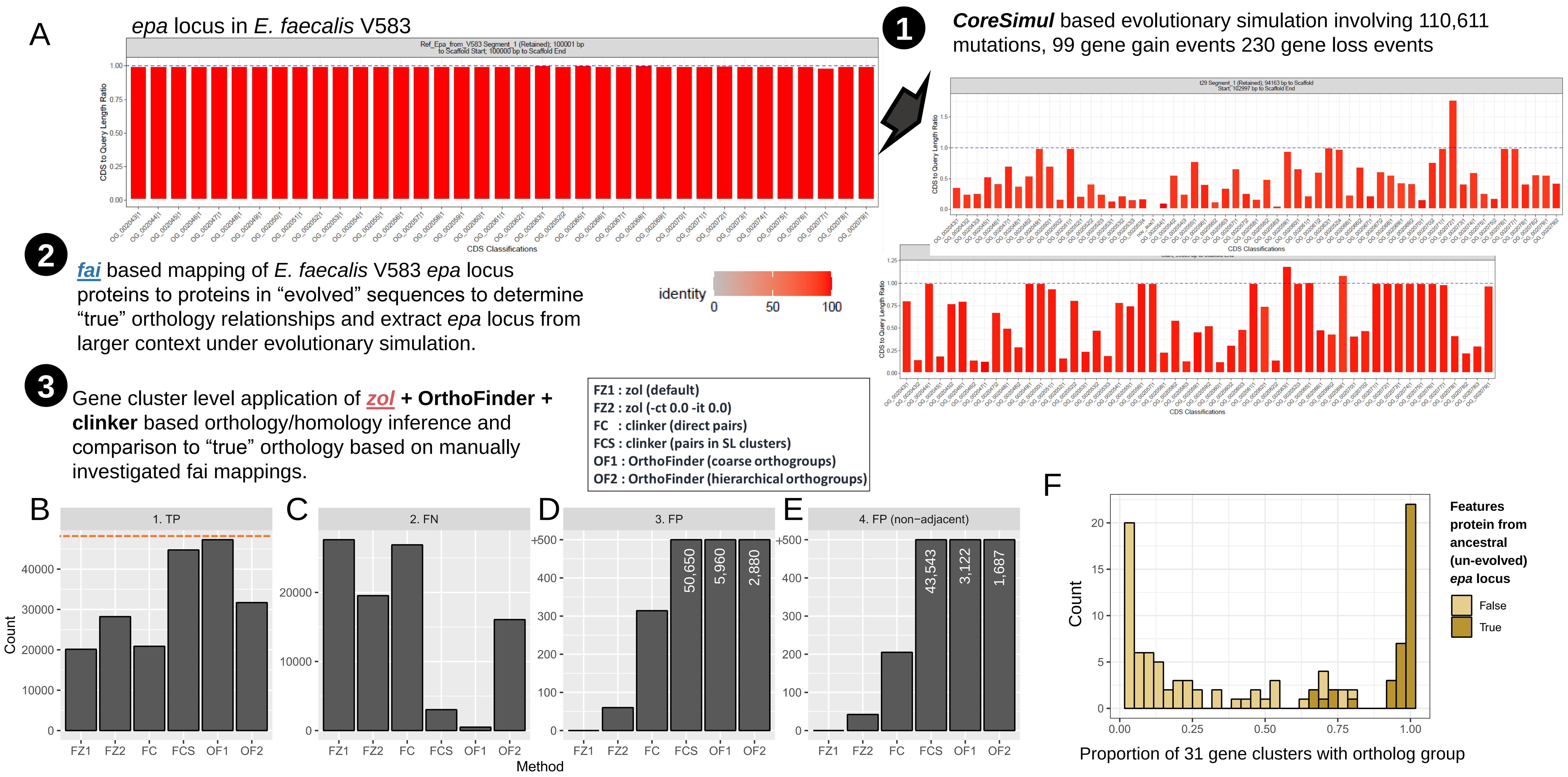


**Figure S7: Overview of simulation-based assessment of ortholog group inferences by zol. A**) An overview of the evolutionary simulation-based benchmarking for context-limited orthology inference. Simulation of the *epa* locus and surrounding +/-100 kb context from *E. faecalis* V583 was performed using CoreSimul to yield 30 artificially evolved sequences. These sequences were then processed, along with the original unevolved sequence, through prepTG to perform gene calling and the core *epa* locus extracted using fai. Visual inspection of figures from fai highlighting homology to *epa* proteins used for determining the locus in sequences was used to determine how truncated or expanded coding sequences should be grouped together with regards to the original query proteins. Coding sequences from regions extracted from the 31 sequences were investigated for orthology/homology using zol, clinker, and OrthoFinder. Comparison of orthology predictions to the inferred truth set were performed to determine **B**) true positives (TP) [dashed red line indicates the number of true positive ortholog pairs], **C**) false negatives (FN), **D**) false positives (FP), and **E**) false positives discounting orthology predictions for neighboring genes (FP (non-adjacent)). **F**) A histogram showing the proportion of gene clusters that feature each ortholog group identified by zol with loosened thresholds for orthology consideration (FZ2; -ct 0.0 -it 0.0). The coloring indicates whether ortholog groups feature a protein sequence from the ancestral instance of the gene cluster.

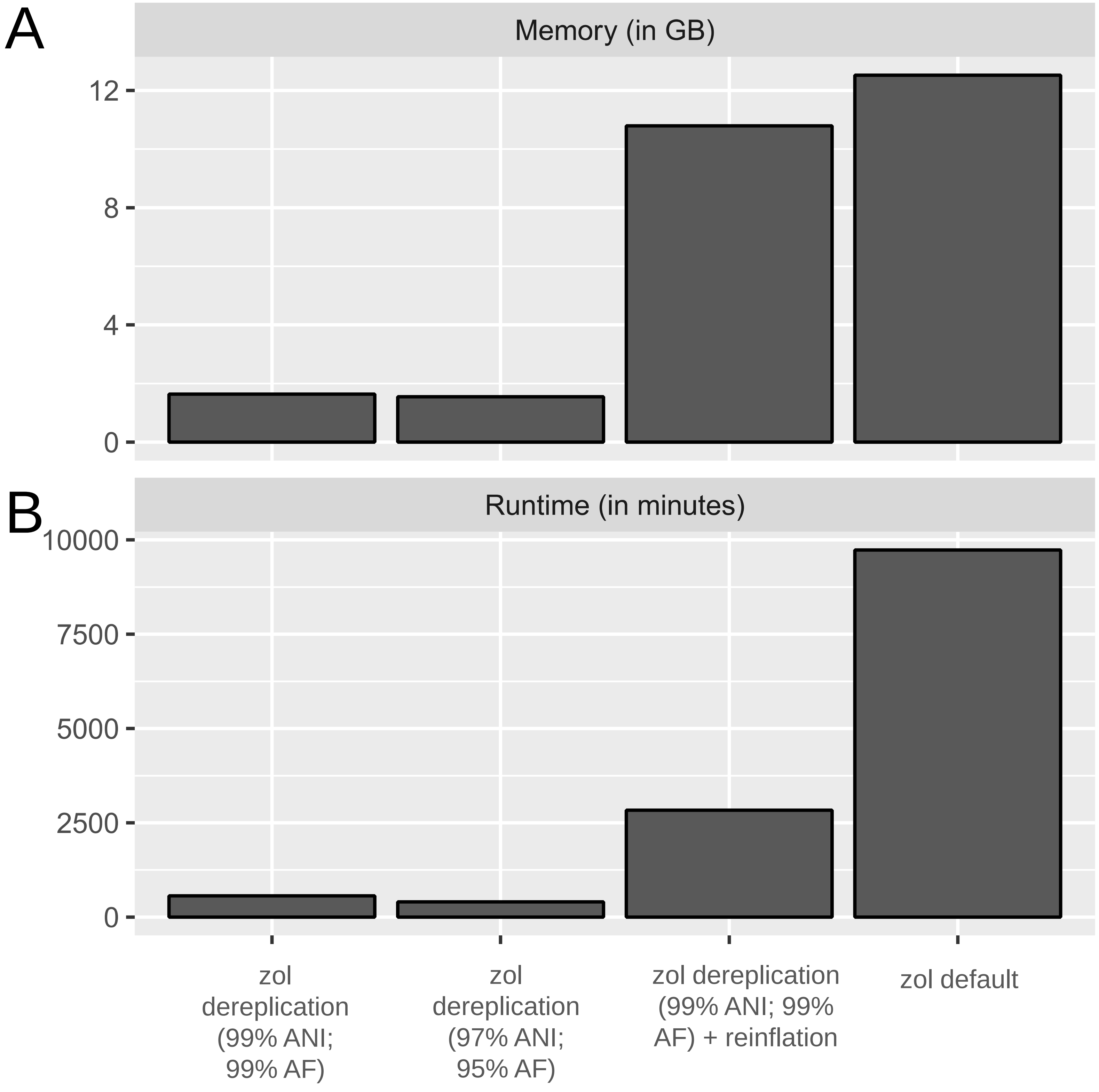


**Figure S8: Time and memory differences for investigation of 1,232 *E. faecalis epa* loci using different approaches within zol. A**) Maximum memory usage and **B**) time to completion for running zol analysis on 1,232 *E. faecalis epa* loci using different options showcasing the ability of dereplication and reinflation to alleviate computational resources and time needed to process large sets of gene-clusters.

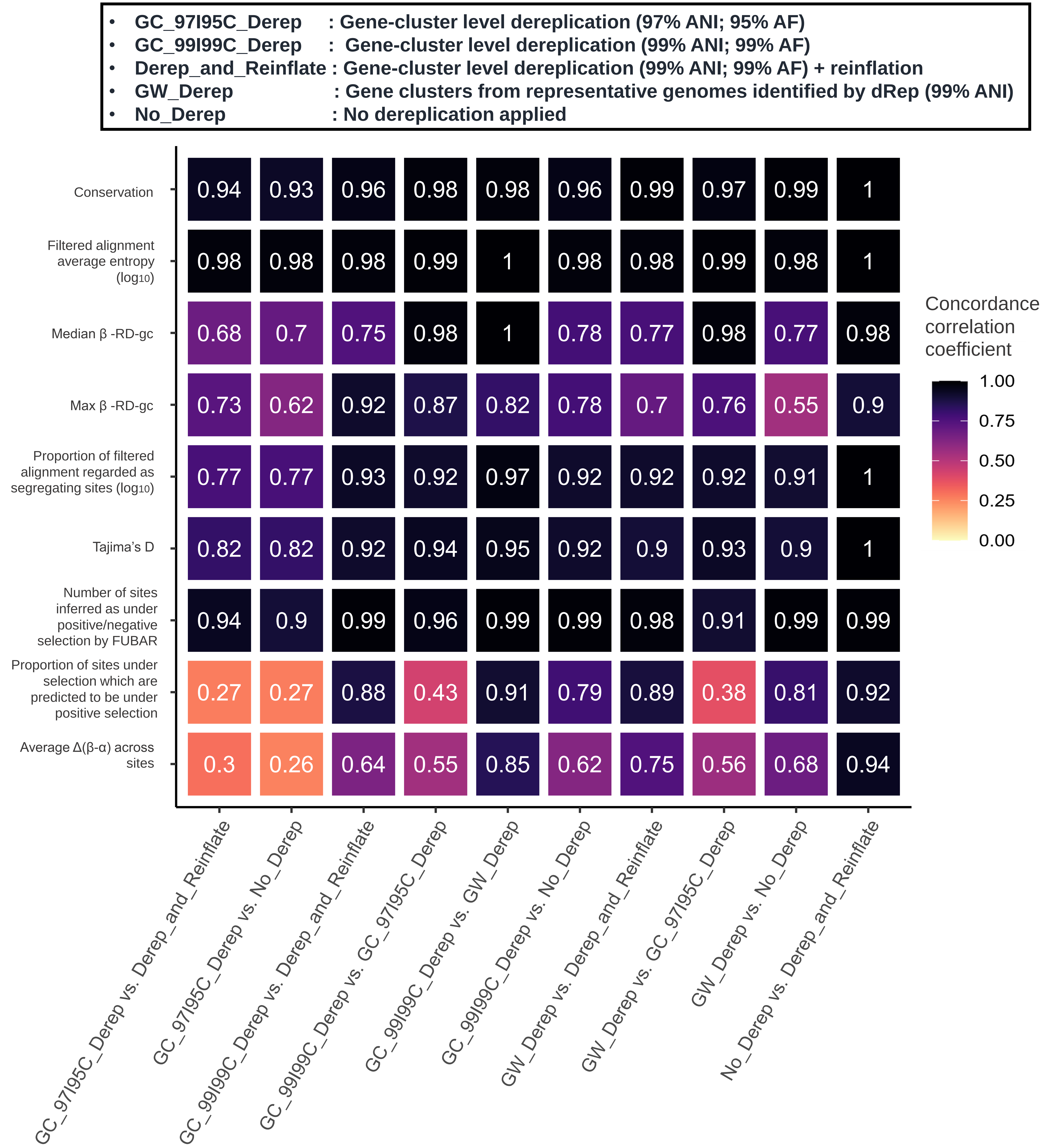


**Figure S9: Effects of dereplication on the calculation of evolutionary statistics by zol.** The heatmap shows the Concordance correlation coefficient for analogous ortholog groups of the *epa* locus for various evolutionary statistics computed by zol when different approaches to dereplication are used.

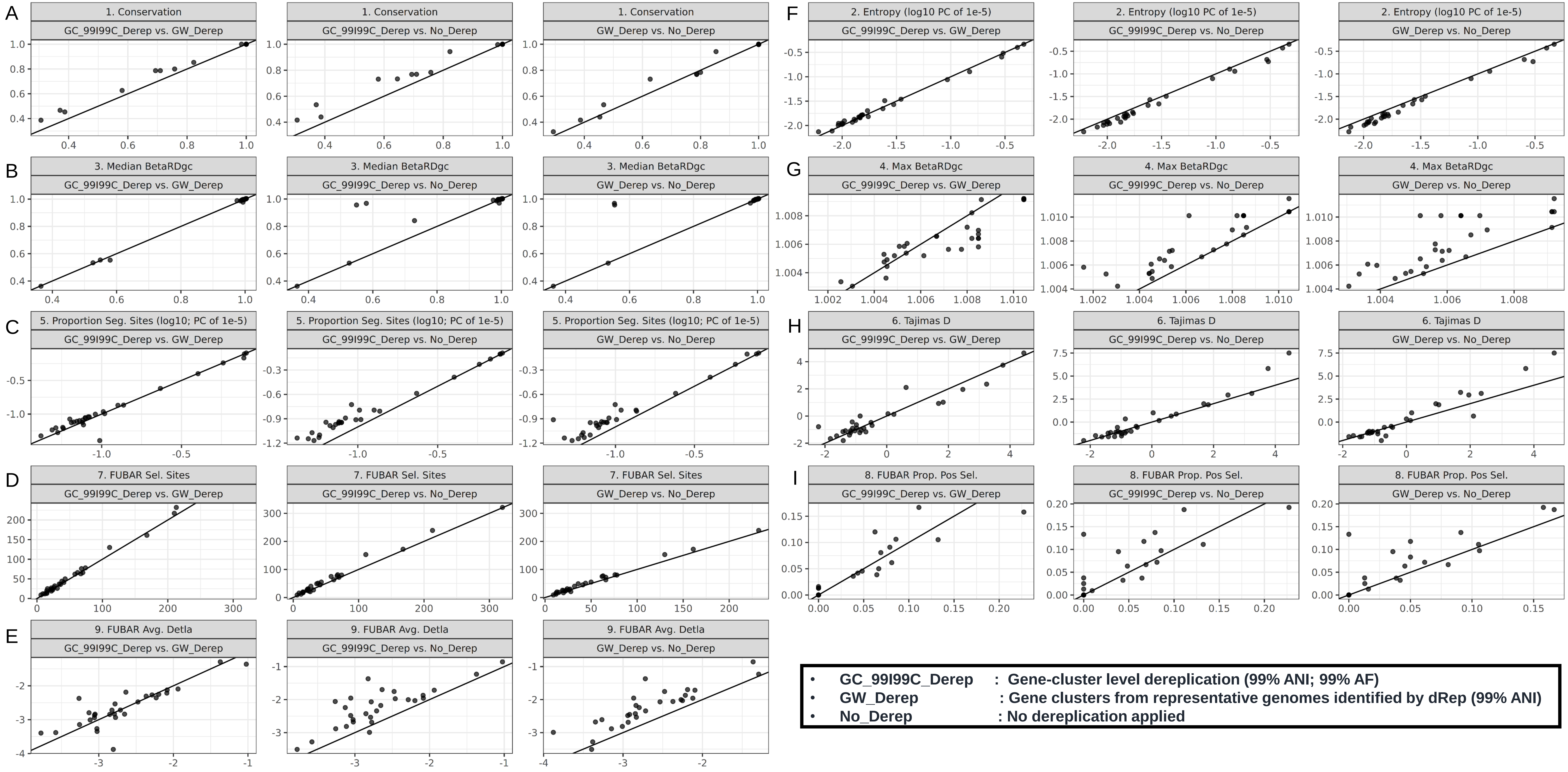


**Figure S10: Influence of dereplication on evolutionary statistics computed by zol**. The relationship in values for analogous ortholog groups which map to query proteins from *E. faecalis* V583 for different evolutionary statistics (**A-I**) when different sets of gene-clusters corresponding to different approaches in dereplication are shown. Only ortholog groups which lacked any paralogous proteins are shown and accounted for. A line is shown in each plot corresponding to a 1:1 ratio.
